# Defining and targeting adaptations to oncogenic KRAS^G12C^ inhibition using quantitative temporal proteomics

**DOI:** 10.1101/769703

**Authors:** Naiara Santana-Codina, Amrita Singh Chandhoke, Qijia Yu, Beata Małachowska, Miljan Kuljanin, Ajami Gikandi, Marcin Stańczak, Sebastian Gableske, Mark P. Jedrychowski, David A. Scott, Andrew J. Aguirre, Wojciech Fendler, Nathanael S. Gray, Joseph D. Mancias

## Abstract

Covalent inhibitors of the KRAS^G12C^ oncoprotein have recently been developed and are being evaluated in clinical trials. Resistance to targeted therapies is common and likely to limit long-term efficacy of KRAS inhibitors (KRASi). To identify pathways of adaptation to KRASi and to predict drug combinations that circumvent resistance, we used a mass spectrometry-based quantitative temporal proteomics and bioinformatics workflow to profile the temporal proteomic response to KRAS^G12C^ inhibition in pancreatic and lung cancer 2D and 3D cellular models. We quantified 10,805 proteins across our datasets, representing the most comprehensive KRASi proteomics effort to date. Our data reveal common mechanisms of acute and long-term response between KRAS^G12C^-driven tumors. To facilitate discovery in the cancer biology community, we generated an interactive ‘KRASi proteome’ website (https://manciaslab.shinyapps.io/KRASi/). Based on these proteomic data, we identified potent combinations of KRASi with PI3K, HSP90, CDK4/6, and SHP2 inhibitors, in some instances converting a cytostatic response to KRASi monotherapy to a cytotoxic response to combination treatment. Overall, using our quantitative temporal proteomics-bioinformatics platform, we have comprehensively characterized the proteomic adaptations to KRASi and identified combinatorial regimens to induce cytotoxicity with potential therapeutic utility.

## INTRODUCTION

*KRAS* is the most frequently mutated oncogene in tumors with 20% of all solid tumors containing oncogenic *KRAS* mutations (Singh et al., 2009) that drive tumorigenesis and tumor maintenance (Cox et al., 2014; Kimmelman, 2015). Chief among these tumors are non-small cell lung cancer (NSCLC) and pancreatic ductal adenocarcinoma (PDAC). Patients with these tumors have dismal survival rates and few available effective therapies; therefore, targeting oncogenic KRAS is a major priority for these patients and the clinicians that treat them. Until recently, efforts to pharmacologically target mutant KRAS directly have been unsuccessful, suggesting that oncogenic KRAS is “undruggable.” However, development of clinical grade KRAS^G12C^ inhibitors has now ushered in a new era of clinical trials testing the efficacy of direct pharmacologic inhibition of KRAS^G12C^ in patients with tumors harboring oncogenic KRAS^G12C^ driver mutations with early promising anti-tumor activity reported in Phase I trials (NCT03600883) (Fakih et al., 2019). A critical lesson from prior clinical experience targeting driver oncoproteins, such as HER2, EGFR, and BRAF, is the development of therapeutic drug resistance that limits efficacy (Piotrowska et al., 2015; Takezawa et al., 2012). Drug resistance to targeted therapy can be multifactorial and heterogenous between patients with the same tumor type and between different tumor types driven by the same oncogene (Konieczkowski et al., 2018).

Identifying mechanisms of drug resistance is a challenge with no one method capable of identifying mutational adaptations or non-mutational adaptations, such as alterations in protein levels, signaling pathways, or metabolic changes, that can all drive cancer cell survival despite inhibition of the main oncogenic driver protein. Successful identification of mechanisms of drug resistance in patients or in pre-clinical studies has led to the development of useful strategies to prevent or circumvent resistance, such as next-generation targeted inhibitors that mitigate development of resistance or combination therapy that circumvents resistance (Bryant et al., 2019; Janne et al., 2015). Initial efforts at defining resistance to KRAS inhibition and identifying synthetic lethal interactions specific to oncogenic KRAS-driven cancers relied on models using genetic ablation of oncogenic KRAS, a situation distinct from pharmacological inhibition. For example, several genetic ablation studies have described how KRAS targeting leads to compensatory mechanisms (Chen et al., 2018; Kapoor et al., 2014; Muzumdar et al., 2017; Santana-Codina et al., 2018; Viale et al., 2014; Ying et al., 2012). More recent efforts with pharmacologic KRAS^G12C^ inhibitors have included targeted investigation of specific collateral signaling pathways (Lito et al., 2016; Misale et al., 2018) and genome-wide combination KRASi with CRISPR/Cas9 screens to identify collateral dependencies (Lou et al., 2019).

Our recent effort using multiplexed mass spectrometry-based quantitative proteomics to identify and target resistance to glutaminase inhibition (GLSi) in PDAC (Biancur et al., 2017) suggests an additional method for investigating the collateral dependencies or adaptations to pharmacologic KRAS^G12C^ inhibition. Using this approach, we identified enriched pathways and targeted these pathways in combination with GLSi demonstrating an effective combinatorial approach with GLSi and glutathione metabolism inhibition with buthionine sulfoximine (Biancur et al., 2017). The potential advantage of this proteomics-first approach to investigate mechanisms of resistance is the identification of specific upregulated proteins or pathways based on direct measurement of the levels of proteins, which are the ultimate target of most therapeutic agents.

Here, we applied our mass spectrometry-based quantitative proteomics to profile the temporal proteomic response to KRAS^G12C^ inhibition in cellular models of KRAS^G12C^-driven PDAC and NSCLC with the ultimate goal of leveraging these results to identify useful combinatorial approaches that prevent or circumvent resistance. We identified and quantified 10,805 proteins across our proteomic datasets allowing us to gain a comprehensive understanding of the global proteomic response to KRASi. Based on these KRASi proteomic signatures, we identified candidate combination therapies to further inhibit pathways supported by KRAS or combination therapies to counteract compensatory survival pathways. We identified multiple combination therapies that prevented re-establishment of proliferation seen in long-term KRASi and in some instances converted cytostatic responses to cytotoxicity.

Our work establishes a proteomic and bioinformatic workflow for identifying the proteomic adaptations to oncoprotein inhibition that can be targeted to improve therapeutic efficacy (Figure 1A). Finally, our temporal KRASi proteomic datasets constitute a significant new resource for the community of scientists studying oncogenic KRAS biology. We termed this resource the ‘KRASi proteome’ and developed an interactive website to aid in the exploration and discovery of new oncogenic KRAS biology (https://manciaslab.shinyapps.io/KRASi/).

**Figure 1.**
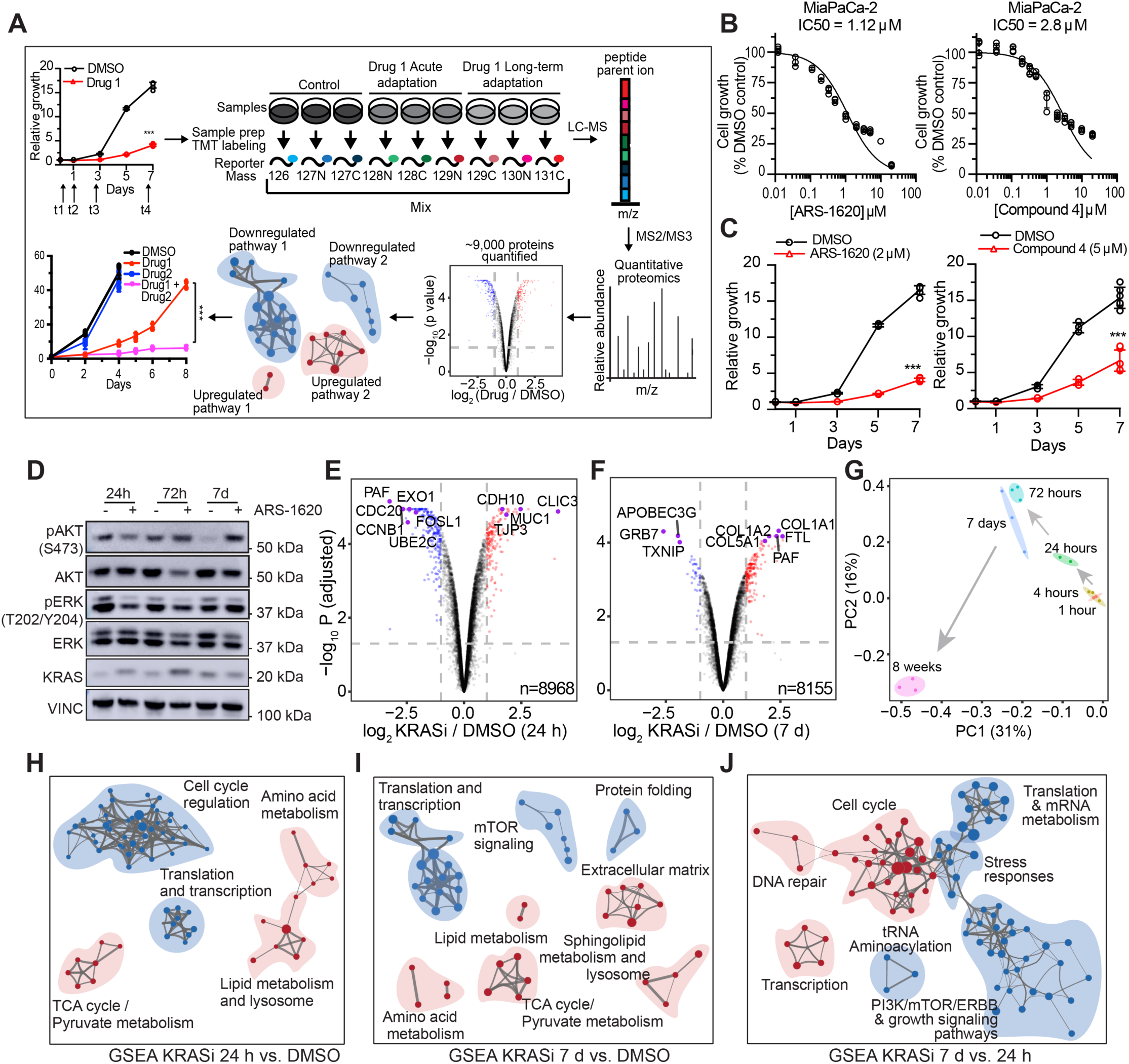
Quantitative temporal KRAS^G12C^ inhibitor proteomics in PDAC cells identifies up- and downregulated pathways. **(A)** Schematic model of the quantitative temporal proteomic-bioinformatic workflow: Cancer cells treated with a targeted inhibitor (Drug 1) at different timepoints are analyzed using multiplexed Tandem Mass Tag (TMT) quantitative proteomics (9 samples shown here, this can now be expanded to 16 samples). Samples labeled with TMT reporter tags are analyzed by LC-MS followed by MS2/MS3 fragmentation for relative quantitation. Volcano plot illustrates protein abundance differences in treated cells. Red and blue: up-and downregulated proteins. Enrichment map of dysregulated pathways. Relative proliferation of cell line treated with Drug 1 in combination with Drug 2 to suppress or circumvent resistance. **(B)** Cell proliferation dose-response curves for MiaPaCa-2 cells treated with KRAS^G12C^ inhibitors (left: ARS-1620, right: Compound 4). Error bars represent s.d. of 4 technical replicates (representative of 3 (ARS-1620) and 5 (Compound 4) independent experiments). **(C)** Relative proliferation of MiaPaCa-2 cells treated with KRASi. Error bars represent s.d. of 6 technical replicates (representative of 3 independent experiments). Significance determined with t-test. ***p<0.001. **(D)** Immunoblot of lysates from MiaPaCa-2 cells treated with ARS-1620 for the indicated times. **(E-F)** Volcano plots illustrate statistically significant protein abundance differences in MiaPaCa-2 cells treated with KRASi for 24 h **(E)** and 7 days **(F)**. Volcano plots display the −log_10_ (P value) versus the log_2_ of the relative protein abundance of mean KRASi treated to mean control (DMSO) samples. Red circles represent significantly (adjusted p value/FDR < 0.05) upregulated proteins (FC ≥ 2) while blue circles represent significantly downregulated proteins (FC < 0.5). Data are from three independent plates from a representative experiment of 2. **(G)** Principal Component Analysis of KRASi treated proteomes (1 h, 4 h, 24 h, 72 h, 7 days and 8 weeks) represented in a two-dimensional space. Ratio of KRASi vs DMSO is used as the data input. **(H-J)**, Enrichment map of gene set enrichment analysis (GSEA) of KRASi-treated MiaPaCa-2 cells versus DMSO at 24 h **(H)** and 7 d **(I)** or 7 d KRASi compared to 24 h KRASi **(J)**. FDR< 0.01, Jaccard coefficient>0.25, node size is related to the number of components identified within a gene set and the width of the line is proportional to the overlap between related gene sets. GSEA terms associated with upregulated (red) and downregulated (blue) proteins are colored accordingly and grouped into nodes with associated terms.

### Experimental Methods

<u>Full details of experimentation are provided in the Supplemental Experimental Procedures</u>

#### Cell Culture

Cell lines were carefully maintained in a centralized cell bank, and routinely inspected for mycoplasma contamination using PCR. All cell lines were maintained at 37°C with 5% CO_2_ and grown in DMEM (Invitrogen 11965) or RPMI 1640 (Invitrogen 11875), supplemented with 10% Fetal Bovine Serum (FBS) and 1% Penicillin/Streptomycin. Cells were grown on standard tissue culture plates for 2D culture or on ultra-low attachment plates for 3D culture.

#### Cell Proliferation Assay

Cells were plated at 3000-5000 cells per well in 24-well plates. Cells were treated as indicated, fixed with 10% formalin, and stained with 0.1% crystal violet. Dye was extracted with 10% acetic acid, and relative proliferation was determined by measuring OD at 595nm.

#### IC50 Assay

Cells were plated in 96-well plates at 2000 cells per well and treated with the indicated compounds. Cell viability was measured after 72 hours, using Cell-Titer Glo (Promega, G7570) assay.

#### Cell death assay

Cells were plated, treated for 72 h with the indicated compounds, stained with Annexin-FITC and propidium iodide (PI) (BD Biosciences 556547), and analyzed using a Beckman Coulter Cytoflex.

#### Western Blot Analysis

Cells were lysed in RIPA buffer. 30-50 µg of protein was resolved on 4-12% SDS-PAGE gels (Thermo Fisher, NP0322) and transferred to polyvinylidene difluoride (PVDF) membranes (Bio-Rad). Membranes were incubated with primary antibodies overnight at 4°C followed by incubation with peroxidase-conjugated secondary antibody. Antibodies used are listed in the Supplemental Experimental Procedures.

#### Metabolomics

Steady state metabolomics experiments were performed as previously described (Son et al., 2013; Sousa et al., 2016).

#### Quantitative Proteomics

Mass spectrometry-based quantitative proteomics were performed as previously described (Biancur et al., 2017; Paulo et al., 2017). Briefly, 50 µg of digested peptides from each sample were labelled with TMT reagent (100 µg) and combined at a 1:1:1:1:1:1:1:1:1:1:1 ratio. Data were obtained using an Orbitrap Fusion Lumos mass spectrometer (Thermo Fisher Scientific, San Jose, CA, USA) coupled with a Proxeon EASY-nLC 1200 LC pump (Thermo Fisher Scientific). Mass spectra were processed using a Sequest-based-in-house software pipeline as described previously (Paulo et al., 2015).

#### Bioinformatic Analysis

Quantitative proteomics data was analyzed with the LIMMA software package (3.40.2) (Ritchie et al., 2015) in R. Pairwise comparisons were performed among experimental conditions, and p values were adjusted through the Benjamini-Hochberg method, which is referred to as false discovery rate (FDR). Gene set enrichment analysis (GSEA) of each comparison was performed with Broad GSEA software (3.0) (Subramanian et al., 2005) using the collection containing all canonical pathways (c2.cp.v6.2.symbols.gmt) in MSigDB for pathway annotation. Cytoscape 3.7.0 software was used to generate Enrichment maps. To retrieve compound inhibitors with similar expression signatures, the top 150 up- and down-regulated proteins were selected and queried against the L1000 gene expression database in Connectivity Map (CMap) 2.0 (Subramanian et al., 2017). Meta-analysis of pathways and protein-protein interaction (PPI) enrichments were performed using Metascape (Zhou et al., 2019).

#### Chemicals

ARS-1620 (Chemgood, C-1454), Compound 4 (Zeng et al., 2017), 17-AAG (Cayman chemicals 75747-14-7), GDC-0941 (Selleckchem, S1065), (PF 2341066 (crizotinib, Cayman chemicals, 877399-52-5), Erlotinib hydrochloride (Cayman chemicals, 183321-74-6), CINK4 (CDK4/6 inhibitor, Cayman chemicals, 359886-84-3), SHP099 hydrochloride (SHP2i, Selleckchem, S8278), CHIR-99021 (GSK3α/βi, Selleckchem S1263).

#### Statistics

All data except the proteomics (analysis described above) was analyzed using GraphPad PRISM software. No statistical methods were used to predetermine sample size. For comparison between two groups, Student’s t-test (unpaired, two-tailed) was performed for all experiments. The level of statistical significance was declared at 0.05 for all analyses.

## RESULTS

### Defining mechanisms of adaptation to KRASi in PDAC by quantitative proteomics

To elucidate mechanisms of adaptation to pharmacologic KRASi, we initially tested the sensitivity of a KRAS^G12C^ mutant pancreatic ductal adenocarcinoma (PDAC) cell line (MiaPaCa-2) to two compounds that selectively target KRAS^G12C^: ARS-1620 (Janes et al., 2018; Misale et al., 2018) and ‘Compound 4’ (Zeng et al., 2017). IC50 measurements (Figure 1B) were in the range of previous studies of this cell line with ARS-1620 (Janes et al., 2018). Both KRASi compounds significantly reduced growth of MiaPaCa-2 cells (Figure 1C), although cells treated with IC50 doses were able to re-establish proliferation at later time points (7 days). KRAS mediates activation of several downstream pathways, with the MAPK pathway being the main mediator of its tumorigenic effects in PDAC (Collisson et al., 2012; Santana-Codina et al., 2018; Ying et al., 2012). To confirm efficient block of KRAS activity, we assessed MAPK and PI3K pathways by western blot (Figure 1D). KRASi induced an upward electrophoretic shift in the KRAS band as a consequence of covalent binding of the inhibitor to KRAS (Zeng et al., 2017), confirming target engagement. ERK1/2 phosphorylation (Thr202/Tyr204) was decreased, consistent with downregulation of the MAPK pathway (Figure 1D). We noted a progressive increase in AKT phosphorylation with long-term KRASi suggesting compensatory activation of the PI3K pathway (Figure 1D) (Alagesan et al., 2015; Muzumdar et al., 2017; Santana-Codina et al., 2018).

To define the global proteomic response to KRASi, we used multiplexed isobaric tag-based quantitative mass spectrometry (Biancur et al., 2017; McAlister et al., 2014). We compared proteomic changes induced by KRASi in MiaPaCa-2 cells at 1 h, 4 h, 24 h, 72 h, 7 days, and 8 weeks (Figure 1E-F, Figure S1A, Table S1). The magnitude of changes at 1 h and 4 h were less pronounced than those observed at 24 h and longer time points, likely related to the lag in alterations in protein expression after mRNA level changes (Figure 1E, Figure S1A). Indeed, Principal Component Analysis (Figure 1G) showed a shift in PC1 and PC2 components in late (72 h, 7 days, 8 weeks) versus early (1 h, 4 h) time points. Among the most downregulated proteins at the 24 h time point, we found proteins related to cell cycle and DNA repair (CDC20, CCNB1, EXO1) (Figure 1E). Upregulated proteins included those related to adhesion at 24 h (TJP3, CDH10) and 7 days (COL1A1, COL1A2, COL5A1) (Figure 1F).

To gain a more comprehensive understanding of the pathways altered in response to acute versus long-term KRASi, we performed Gene Set Enrichment Analysis (GSEA) on proteomes from KRASi-treated cells at 24 h (acute) and 7 days (long-term) (Figure 1H-J, Table S2). Enrichment map nodes associated with upregulated proteins at 24 h included TCA cycle/pyruvate metabolism, fatty acid/lipid and amino acid metabolism as well as lysosomal processes (Figure 1H). These results are consistent with studies in genetically KRAS-ablated cells that demonstrated a compensatory metabolic response consisting of upregulated oxidative phosphorylation, autophagy, and lipid metabolism (Viale et al., 2014). Enrichment map nodes associated with downregulated proteins included cell cycle and translation/transcription pathways consistent with a decrease in proliferation (Figure 1H). To identify if deeper inhibition of oncogenic KRAS altered the landscape of proteomic changes, we analyzed the proteome after KRASi at an IC90 dose level. In addition to pathways identified at the IC50 dose level, we identified downregulation in DNA damage associated pathways and upregulation of interferon signaling pathways and extracellular matrix (ECM) associated pathways (Figure S1B).

Several of the biological processes upregulated at 24 h were consistently activated at 7 days including TCA cycle, amino acid and lipid metabolism (Figure 1I). Interestingly, long-term KRASi induced down-regulation of mTOR signaling (Figure 1I) and upregulation of terms associated with lysosomal processes, consistent with recent evidence demonstrating ERK inhibition in PDAC activates autophagy thereby identifying a therapeutically useful combination of ERK inhibition and lysosome inhibition with hydroxychloroquine (Bryant et al., 2019). Notably, cell cycle pathways were no longer downregulated in the long-term, which aligns with reactivation of proliferation (Figure 1I). Comparative GSEA analysis of KRASi proteomes at 7 days versus 24 h to identify pathways reversed with long-term inhibition compared to acute inhibition revealed relative upregulation of cell cycle, transcription and DNA repair pathways together with downregulation of stress response signaling pathways (Figure 1J). GSEA of the 8-week KRASi proteome identified upregulation of ECM pathways, fatty acid metabolism and oxidative phosphorylation (oxphos) and a decrease in glucose metabolism, translation, protein-folding and the PI3K-mTOR pathway (Figure S1C).

Next, we selected the 15 top up and downregulated pathways at 24 h and 7 days and analyzed their temporal response (Figure S1D-H). Among the most downregulated pathways at 24 h, we identified cell cycle and DNA repair which were reactivated at longer time-points (Figure S1D). Downregulated pathways at 7 days included RNA metabolism and translation as well as glucose and pentose phosphate metabolism or actin cytoskeleton (Figure S1E) which demonstrated a progressive downregulation over time. Among the upregulated pathways, TCA cycle, lipid and amino acid metabolism were upregulated at 24 h and maintained activation at 7 days (Figure S1F, G) while sphingolipid metabolism and ECM peaked at longer time-points (Figure S1G). Finally, mTOR downregulation was inversely correlated with a temporal increase in lysosomal pathways (Bryant et al., 2019) (Figure S1H).

Our results suggest that KRASi triggers a compensatory activation of several metabolic pathways. Indeed, oncogenic KRAS induces a metabolic reprogramming that consists of increased flux through glycolysis, the non-oxidative pentose phosphate pathway (PPP) and the hexosamine biosynthesis pathway (HBP) to support glycosylation and DNA synthesis (Santana-Codina et al., 2018; Ying et al., 2012). To analyze the metabolic changes induced by KRAS^G12C^ inhibitors over time, we performed targeted liquid chromatography-tandem mass spectrometry (LC-MS/MS) at 24 h, 72 h and 8 weeks after KRASi treatment. KRASi at 24 h revealed downregulation of glycolysis, non-oxidative PPP and nucleotide metabolism (Figure S2A-D), consistent with our previous studies (Santana-Codina et al., 2018; Ying et al., 2012). Interestingly, despite ongoing KRASi treatment, at long-term time points these pathways were all reactivated (72 h and 8 weeks, Figure S2A-D), suggesting mechanisms to reengage the same metabolic dependencies in order to support proliferation.

To assess phosphorylation-dependent signaling pathways after KRASi, we performed a global quantitative phosphoproteomic analysis identifying 21,048 phospho-peptides (Table S3). KRASi induced minor differences in the 1 h and 4 h phosphoproteome (Figure S3A-B) but significant decreases in phosphorylated proteins after 24 h (Figure S3C-D). Substrates within the KRAS pathway (MAPK1, BRAF) were downregulated as early as 4 h consistent with repressed activation of the pathway. Likewise, FOSL1, a recently described downstream KRAS mediator that is preferentially phosphorylated and activated in KRAS mutant cells, showed a decrease in phosphorylation upon KRAS inhibition (Lou et al., 2019; Vallejo et al., 2017). To identify the overall response pattern, we performed gene ontology (GO) enrichment analysis (biological processes) at 24 h, which revealed inhibition in DNA replication and cell cycle consistent with a cytostatic response (Figure S3E-F). To understand if differences in phosphorylation were driven only by translational changes, we selected the top up and downregulated phospho-peptides and normalized to protein expression filtering by relevance of phosphosite to cancer and therapeutic resistance (Figure S3G). Among the proteins that revealed decreased phosphorylation, we identified phosphosites that have been previously correlated with KRAS signaling in cell cycle checkpoint and DNA repair such as CDC20, RIF1, RB1, CLSCPN and EXO1 (Franco et al., 2016; Stuart et al., 2015) as well as more generally in lung and breast cancer (Kim et al., 2013; Schweppe et al., 2013; Yi et al., 2014). A small number of phospho-peptides were upregulated after KRASi such as sites in adhesion molecules (TNS1, FAT4 and TJP3) (Britton et al., 2014; Mertins et al., 2014), cell cycle repressors (CDKN1B), immune regulators (TRAFD1), and mitochondrial components (NDUFAF3) (Figure S3C, G). Overall, the phosphoproteomics data were in accordance with pathways identified in the global proteomic profile and correlated with a decrease in KRAS signaling that downregulates cell cycle and increases adhesion molecules (Chen et al., 2018).

Together, this work serves as the most comprehensive quantitative temporal proteomic evaluation of KRASi. Our data align with prior targeted studies of KRAS dependencies thereby highlighting the relevance of our dataset for further analysis as demonstrated below and for the KRAS research community. To extend our observations and identify commonalities in KRASi across tumor types, we subsequently profiled KRAS^G12C^ driven lung cancer cell lines.

### Defining mechanisms of adaptation to KRASi in NSCLC by quantitative proteomics

As KRAS^G12C^ is the mutant genotype in 44% of all KRAS mutant NSCLC tumors (Cox et al., 2014), we analyzed proteomic adaptations to KRASi in a KRAS^G12C^ lung cancer cell line, H358 (Figure 2A-B). We confirmed target engagement by upward mobility shift in KRAS on immunoblot (Figure 2C) and pathway block by decreased MAPK signaling at acute time points (24 h). Notably, pERK levels partially recovered at 72 h and 7 day timepoints (Figure 2C) and there were no major differences observed in pAKT activation (Figure 2C).

**Figure 2.**
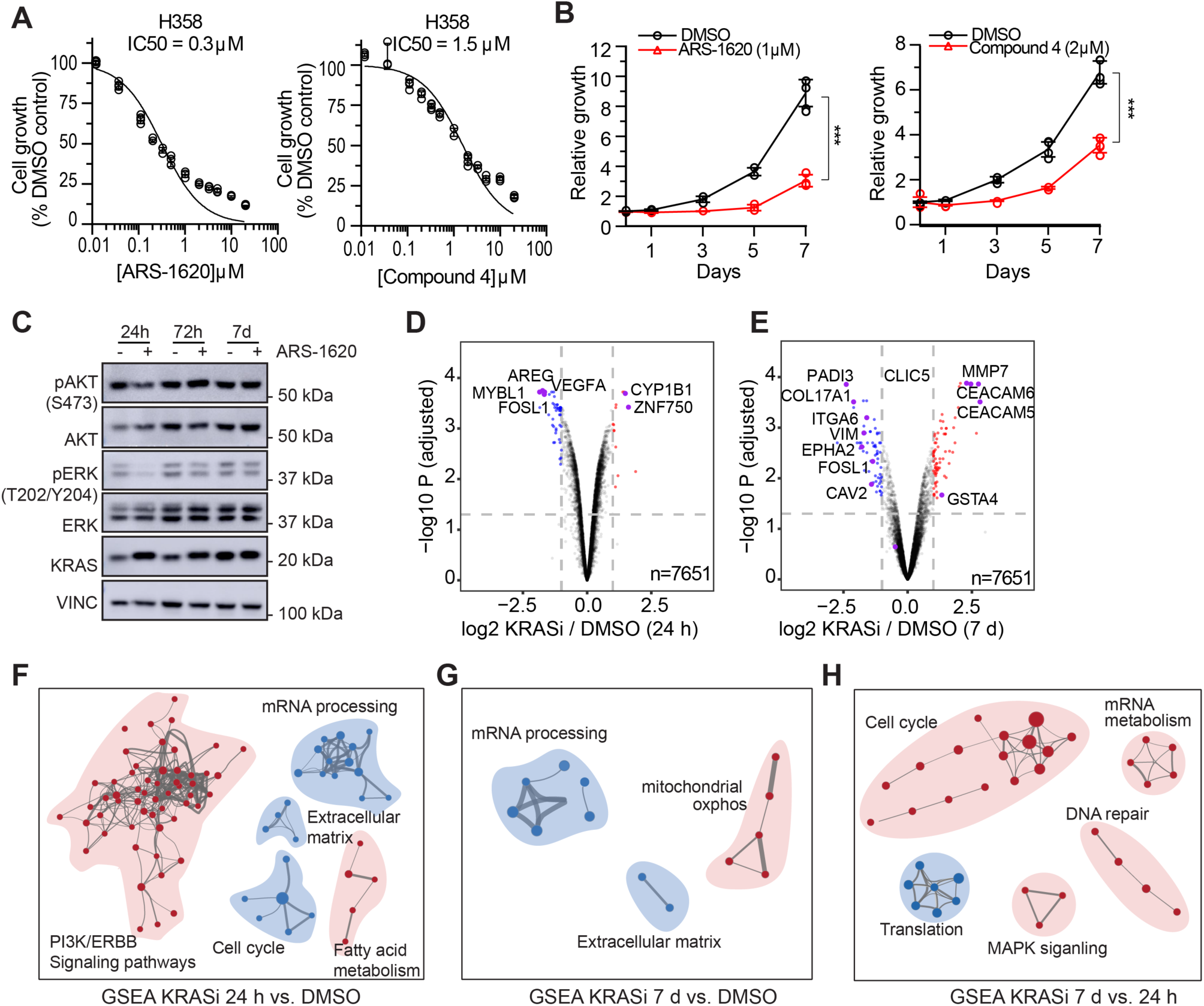
Quantitative temporal KRAS^G12C^ inhibitor proteomics in NSCLC cells identifies up- and downregulated pathways. **(A)** Cell proliferation dose-response curves for H358 cells treated with KRAS^G12C^ inhibitors (left: ARS-1620, right: Compound 4). Error bars represent s.d. of 4 technical replicates (representative of 2 experiments). **(B)** Relative proliferation of H358 cells treated with KRASi. Error bars represent s.d. of 3 technical replicates (representative of 3 independent experiments). Significance determined with t-test. ***p<0.001. **(C)** Immunoblot of lysates from H358 cells treated with ARS-1620 for the indicated times. **(D-E)** Volcano plots illustrate statistically significant protein abundance differences in H358 cells treated with KRASi for 24 h **(D)** and 7 d **(E)** as in Fig. 1E. **(F-H)** Enrichment map of GSEA of KRASi-treated H358 cells versus DMSO at 24 h **(F)** and 7 days **(G)** or 7 d KRASi compared to 24 h KRASi **(H)** as in Fig. 1H-J.

Next, we performed quantitative KRASi proteomic analysis at acute (24 h) and long-term (7 days) time points (Figure 2D-E). GSEA identified similar pathway alterations to those identified in MiaPaCa-2 cells, including upregulation of fatty acid metabolism (24 h, Figure 2F, Table S4) and oxidative phosphorylation (7 days, Figure 2G) and downregulation of cell cycle and mRNA processing. Comparative GSEA of KRASi treated cells at 7 days versus 24 h revealed a relative increase in cell cycle, mRNA metabolism and DNA repair consistent with reactivation of proliferation. Furthermore, the MAPK signaling pathway was upregulated (Figure 2H).

To identify the overlap between MiaPaCa-2 and H358 datasets, we performed a pathway and protein-protein interaction (PPI) enrichment and clustering analysis using the top 300 upregulated proteins at 24 h and 7 days time points using Metascape (Tripathi et al., 2015). Despite minimal specific protein overlap at 24 h (Figure 3A) and 7 days (Figure 3D) between the two cell lines, we observed considerable overlap in the GO terms associated with altered protein expression, suggesting a convergence of pathways in the set of upregulated proteins. Analysis of overlapping functions between cell lines at acute timepoints (24 h) included pathways like metabolism (pentose phosphate and lipid metabolism), cytokine signaling and activation of the PI3K/AKT signaling pathway (Figure 3B-C), while long-term adaptation was associated with alterations in antigen presentation, response to oxidative stress, and lysosomal pathways (Figure 3E-F), in accordance with a general metabolic rewiring that likely involves autophagy and a shift toward mitochondrial metabolism. To identify PPI clusters at 24 h and 7 days, we applied the Molecular Complex Detection algorithm (MCODE, see Methods). We identified PPI clusters involved in signaling pathways including the RAS-MAPK and PI3K/AKT pathways, glycolysis and PPP (Figure 3G) and PPIs associated with cell cycle and ECM in long-term KRASi (Figure 3H). The same analysis for the top 300 downregulated proteins revealed pathways related to cell cycle, DNA repair and transcription as commonly decreased in the acute setting (data not shown).

**Figure 3.**
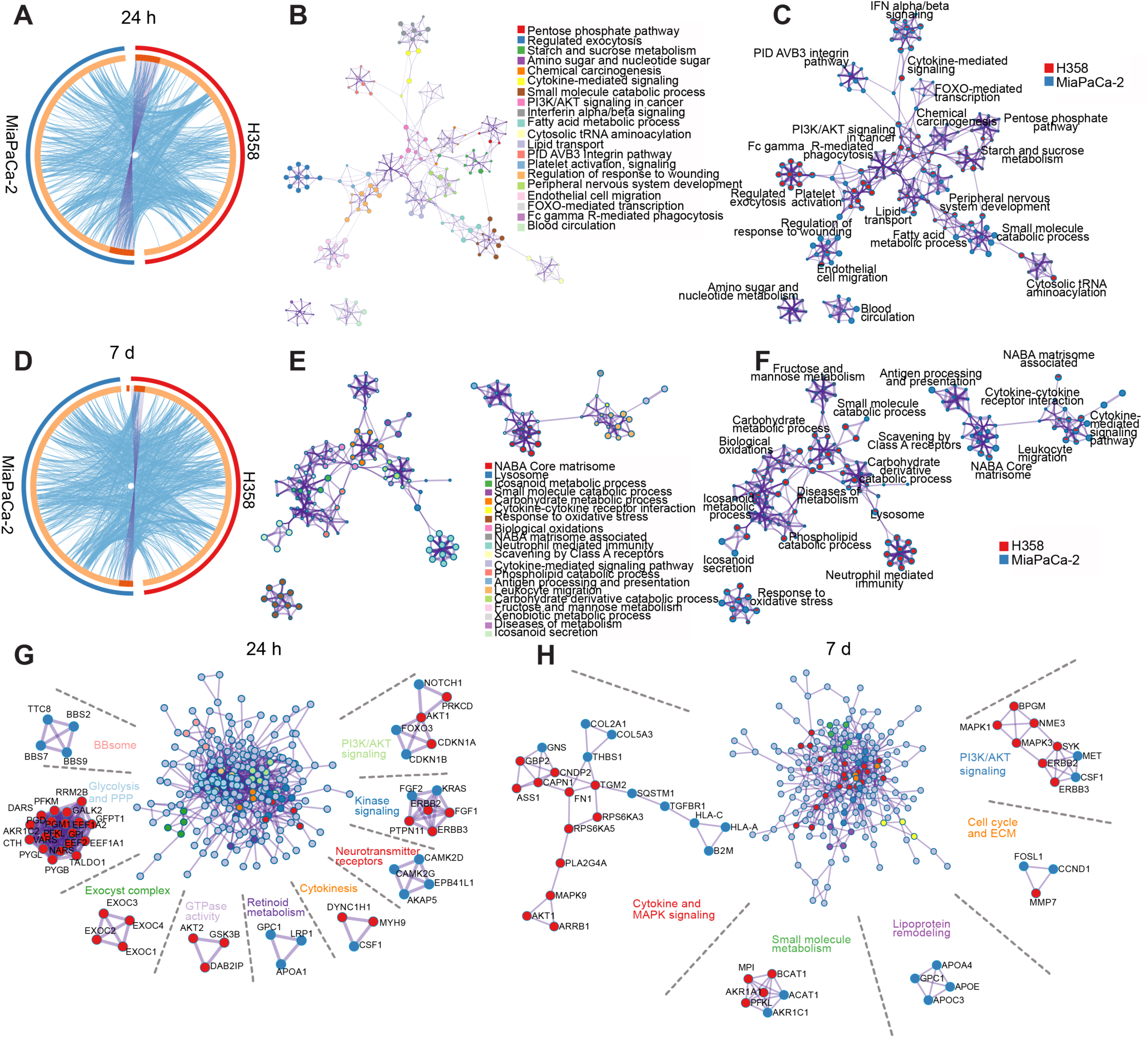
KRASi proteomes from distinct cancer types show similar pathway enrichment. **(A and D)**, Circos plots (see methods) showing overlap of top 300 upregulated proteins and associated gene ontology (GO) terms between MiaPaCa-2 and H358 KRASi proteomes at 24 h **(A)** and 7 days **(D).** The outside arc color identifies each cell line by color. The inside arc represents the top 300 proteins from each proteome: dark orange bars represent the proteins shared between datasets with purple lines linking them, light orange bars represent non-overlapping proteins. Blue lines connect proteins from the same GO term. **(B and E)**, Network shows pathway enrichment at 24 h **(B)** and 7 days **(E).** Each term is identified by color and size of clusters is proportional to the number of input proteins included in the term. **(C and F)** Network of enriched terms represented as pies show contribution of each input list to each term at 24 h **(C)** and 7 d **(F). (G, H)** Protein-protein interaction analysis of input proteins. The center node represents the full interactome and the surrounding nodes are identified according to the representative functions included. Each node is colored according to cell line contribution (red, H358 and blue, MiaPaCa-2).

We were further interested in correlating our proteome data to recent large scale efforts identifying KRAS collateral dependencies in the context of pharmacologic KRAS^G12C^ inhibition. We compared our H358 KRASi proteome datasets (24 h and 7 days) to a recent CRISPR interference (CRISPRi) functional genomic screen with KRASi (Lou et al., 2019) (Figure S4A-B). While this analysis showed poor correlation between datasets on a gene by protein basis, likely given the differential biological and technical approaches used, overall, both analyses identified dependencies in pathways such as cell cycle, cytoskeleton/cell adhesion and lipid biosynthesis (Lou et al., 2019). This suggests that proteome analysis and combination CRISPR/Cas9 screens can provide complementary outputs towards a similar goal of identifying compensatory pathways to target for effective combination therapy. To investigate the correlation between proteome and transcriptome alterations in response to pharmacologic KRASi, we compared MiaPaCa-2 and H358 datasets (KRASi 24 h) with transcriptome profiling in H358 cells treated with KRASi for 24 h (Janes et al., 2018). We observed a positive correlation between these datasets as shown by clustering analysis and comparison of relevant GSEA hallmarks, including ERK signatures and KRAS dependency genes (Janes et al., 2018) (Figure S4C-D).

Our H358 and MiaPaCa-2 proteomics data suggests there are shared proteomic responses to KRASi between different tumor types. To understand whether similarly cytostatic targeted agents produce a conserved proteomic signature, we compared our MiaPaCa-2 KRASi proteomic datasets to a GLSi proteomics analysis performed in a different PDAC cellular model (Biancur et al., 2017). Comparative GSEA analysis revealed a number of shared proteomic responses (Figure S4E-H). Shared pathways associated with downregulated proteins at 24 h and 72 h included cell cycle and translation consistent with a cytostatic response. Furthermore, protein folding pathways were downregulated in both KRASi and GLSi proteomes consistent with upregulation of the endoplasmic reticulum (ER) stress pathways (Biancur et al., 2017; Genovese et al., 2017). Interestingly, fatty acid metabolism and TCA cycle were similarly upregulated at 72 h and long-term, suggesting that activation of oxidative pathways may be a common mechanism of response to cytostatic drugs. Proteins associated with lysosomal processes were upregulated at 24 h and long-term, suggesting autophagy activation as a common response to acute and long-term inhibition with cytostatic drugs. Overall, these results suggest that drugs that induce cytostasis may induce shared mechanisms of proteomic adaptation. Therefore, targeting of these adaptive pathways may be an effective method to overcome resistance induced by any cytostatic drug. Indeed, several of the combination therapies explored below were also effective in combination with GLSi, e.g. ER stress induction (Biancur et al., 2017).

### Targeting adaptation to KRASi

To further understand the functional response to KRASi and identify drug combinations that could bypass or mitigate KRASi resistance, we used the Connectivity Map (CMap version 2.0) (Subramanian et al., 2017) database which contains expression profiles in response to 19,811 compounds and 7,494 genetic perturbations (Table S5). By identifying the compounds that induce a pattern of response positively or negatively correlated to the KRASi proteome, these compounds can be tested in combination with KRASi. The hypotheses are that 1) pushing cells further down a response pathway with a positively correlated compound may lead to cytotoxicity and 2) reversing a proteomic adaptive response by treating with a compound negatively correlated with the KRASi proteome may reverse adaptation to induce cytotoxicity. This strategy has been successfully applied in the setting of GLSi (Biancur et al., 2017). We initially interrogated the MiaPaCa-2 KRASi proteome at 24 h and identified 30 perturbagen classes that positively correlated (CMap > 90) (Figure 4A). Among these perturbagen classes, we found drugs that target downstream elements in the KRAS pathway, such as MEK and RAF inhibitors (Figure 4A). We also identified drugs that have been used in combination with KRAS^G12C^ inhibitors, such as PI3K inhibitors (Engelman et al., 2008; Misale et al., 2018) and those used with KRAS-MAPK pathway inhibitors, including HSP90 (Acquaviva et al., 2012), SRC (Rao et al., 2018), HDAC (Ischenko et al., 2015), IGF1 (Molina-Arcas et al., 2013), mTOR (Fritsche-Guenther et al., 2018), BRAF (Dietrich et al., 2018) and PDGFR/Kit inhibitors (Sahu et al., 2017) (Figure 4A). Connectivity Map analysis also revealed inhibitors of pathways identified by KRASi GSEA, such as cell cycle (CDKi, DNA synthesis inhibitor) and an inverse correlation with tubulin inhibitors, consistent with altered extracellular matrix noted on GSEA and an increased resistance to actin polymerization inhibitors in KRAS-pathway inhibited cells (Chen et al., 2018). Finally, we identified inhibitors with no prior KRASi GSEA association including VEGFR, EGFR, topoisomerase, ribonucleotide reductase and IMPDH pathway inhibition (Figure 4A). These data show that the proteomic profile of KRAS^G12C^ inhibition is similar to the profile of KRAS (WT/mutant)-MAPK signaling pathway inhibition and more broadly correlates with the profile of inhibitors that can similarly induce cytostasis. Furthermore, several of these highly correlated compounds have been previously described as useful combinatorial approaches with KRAS pathway inhibition suggesting that our approach may identify further high priority combinations for evaluation.

**Figure 4.**
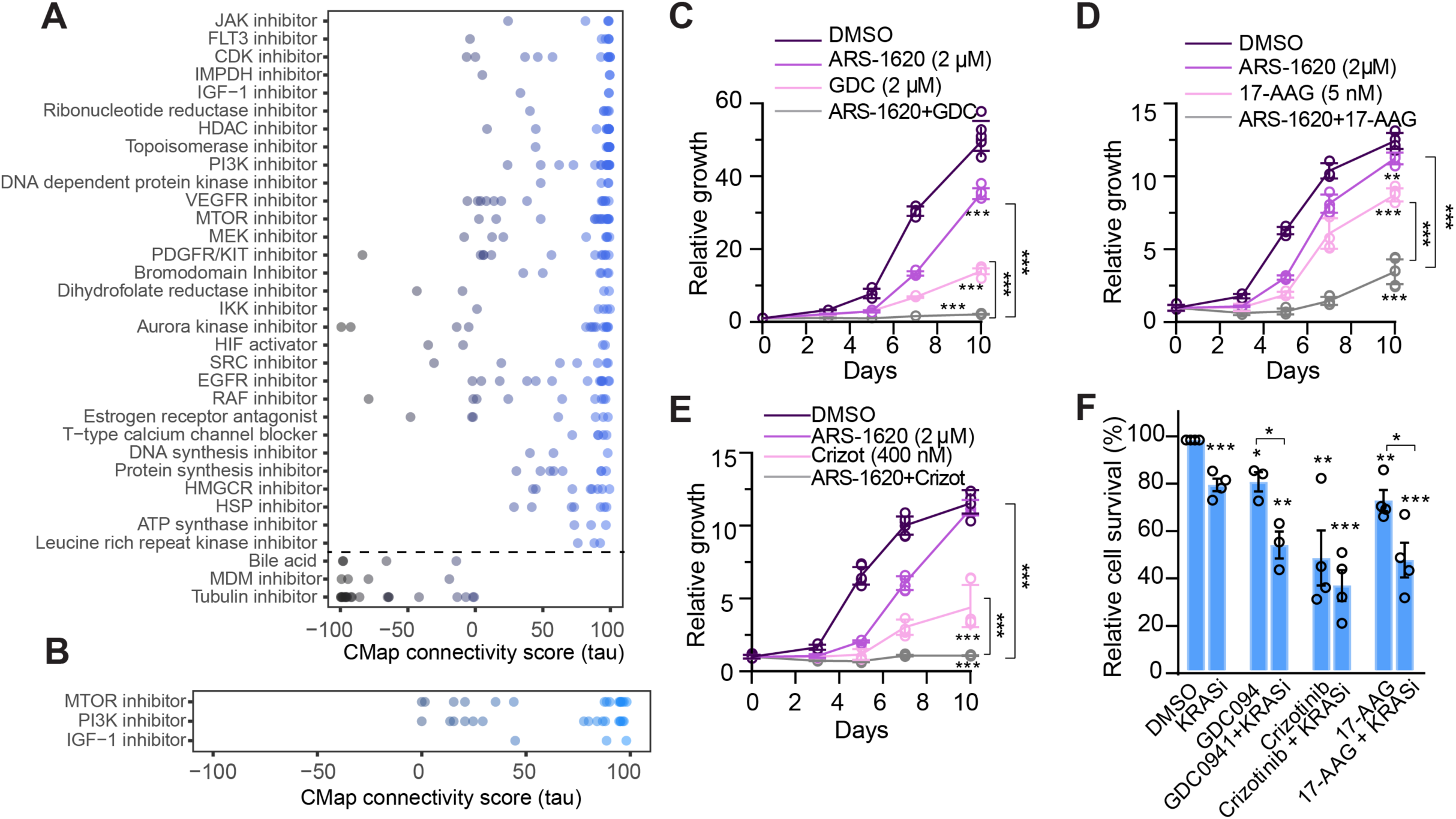
Design of combinatorial pharmacological strategies to overcome adaptation to KRASi. **(A and B)**, Connectivity map analysis for 24 h (**A)** and 7 d **(B)** KRASi MiaPaCa-2 proteomes. Perturbagen classes with mean connectivity scores >90% or <-90% and FDR<0.05 are displayed. **(C-E)** Relative proliferation of MiaPaCa-2 cells treated with KRASi (2 μM) or DMSO with or without GDC0941 (2 μM) **(C)**, 17AAG (5 nM) **(D)** or crizotinib (400 nM). **(F)** Annexin V-FITC and propidium iodide-based flow cytometry analysis of cell death in MiaPaCa-2 cells treated with DMSO/KRASi (2 μM) together with GDC0941 (2 μM), crizotinib (1 μM) or 17-AAG (30 nM) (average of 2 or 3 experiments as indicated). Significance determined with t-test. All comparisons to DMSO except when indicated showing comparison to each drug. *p<0.05, **p<0.01, ***p<0.001.

We also performed this combinatorial proteomic CMap analysis using the 7-day KRASi proteome. Here, we identified a smaller number of positively correlated drugs, namely mTOR, PI3K and IGF1 inhibitors (Figure 4B). These results align with re-establishment of proliferative capacity but sustained inhibition of mTOR signaling demonstrated in GSEA analysis (Fig. 1I).

To identify useful KRASi combination therapies, we evaluated combinations of KRASi with drugs identified in the CMap analysis in long-term proliferation assays. KRAS^G12C^ inhibitors in combination with PI3K inhibitors (GDC0941, Figure S5A) were able to inhibit growth long-term (Figure 4C), in agreement with previous studies (Misale et al., 2018; Muzumdar et al., 2017). Combination with a HSP90 inhibitor (17-AAG) inhibited long-term proliferation (Figure 4D).

In addition to the CMap analysis as a method for identifying combinatorial therapies, we examined upregulated proteins for those with FDA-approved drugs (DrugBank Version 5.1.4) and upregulated cell surface proteins for potential future target investigation (Table S6). Interestingly, in MiaPaCa-2 cells several upregulated druggable proteins were involved in pathways identified in GSEA such as signaling (MET, TGFBR1), lipid metabolism (LDLR) and autophagy (CLU) (Table S6). As the MET pathway is known to be increased in KRAS-mutant lung cancer after MAPK inhibition (Kim et al., 2016) and also after treatment with KRASi in our experiments (Figure S5B), we tested a MET inhibitor (crizotinib) in combination with KRASi, which led to a block in proliferation in MiaPaCa-2 cells (Figure 4E). Finally, we identified two upregulated druggable proteins that overlapped between MiaPaCa-2 and H358 cells: BIRC5 (Survivin) (24 h) (Werner et al., 2016) and P2RX4 (7 days).

As KRASi are generally cytostatic, we checked the ability of the evaluated combinations to induce cytotoxicity. Acute combinatorial treatment induced some level of cell death for multiple combinations (Figure 4F). Of note, combination with a mTOR inhibitor, also identified in the CMap analysis, did not induce differential cell death (data not shown).

CMap analysis of KRASi-treated H358 cells revealed an almost complete overlap with pathways obtained for MiaPaCa-2 cells (Figure 5A, B), confirming a common activation of mechanisms that mediate KRASi effects between different tumor types. We tested the potency of these combinations in long-term growth assays with three of the drugs identified as positive correlators by CMap analysis (Figure S5C). PI3K inhibitors (GDC0941, Figure 5C), HSP90 inhibitors (17-AAG, Figure 5D) and EGFR inhibitors (erlotinib, Figure 5E) were able to block regrowth of KRASi treated cells efficiently when used in combination with KRASi. Furthermore, combination with KRASi led to some level of cytotoxicity in all combinations (Figure 5F). These data strengthen the finding that quantitative proteomics followed by GSEA and CMap analysis are able to predict useful therapeutic drug combinations that overcome cytostasis and induce cytotoxicity.

**Figure 5.**
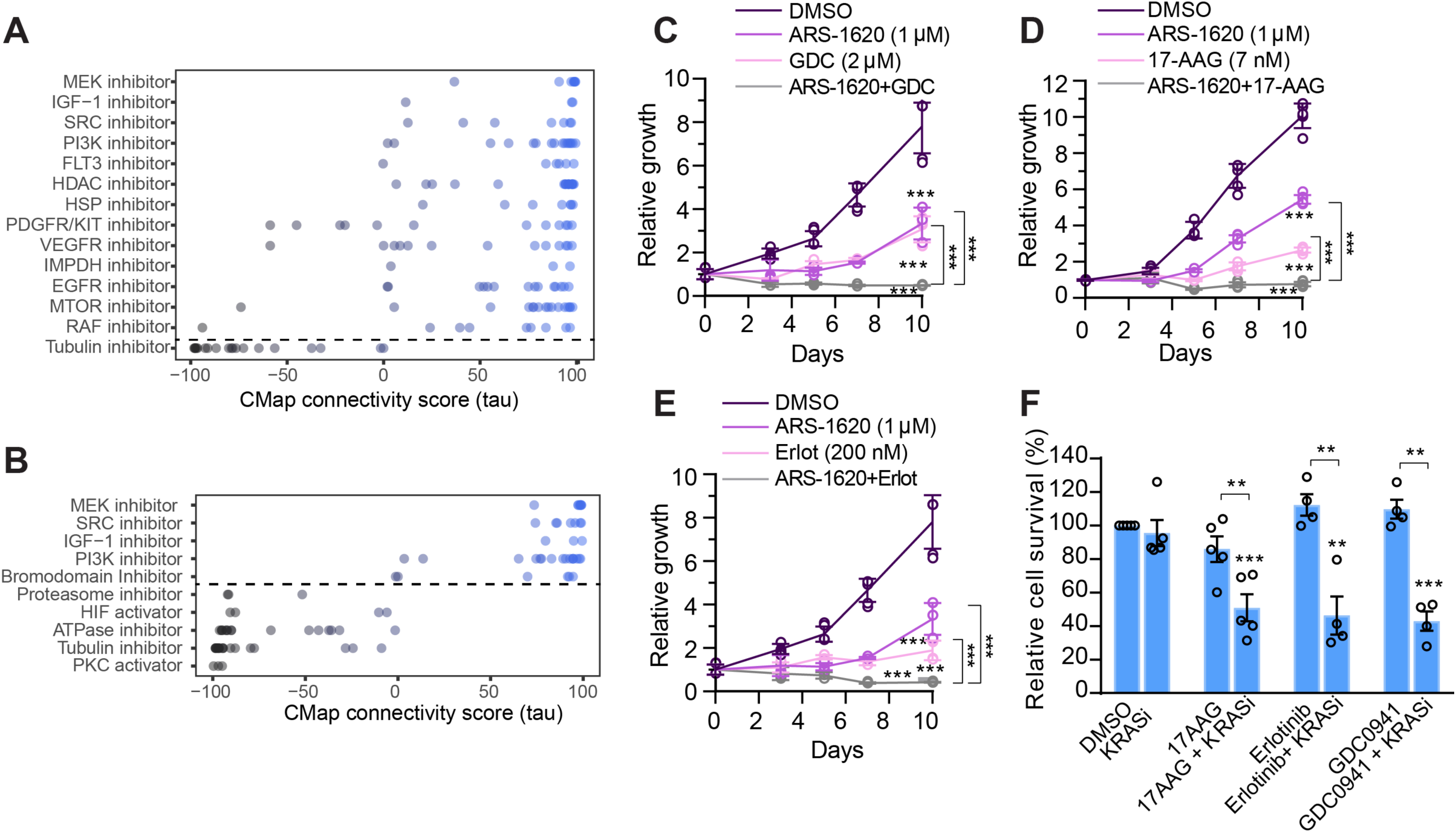
Design of combinatorial pharmacological strategies to overcome adaptation to KRASi in a NSCLC cell line. **(A and B)**, Connectivity map analysis for KRASi H358 proteomes at 24 h (**A)** and 7 days **(B)** as in Fig. 4A. **(C-E)**, Relative proliferation of H358 cells treated with KRASi (1 μM), or DMSO with or without GDC0941 (2 μM) **(C)**, 17-AAG (7 nM) **(D)** and erlotinib (200 nM) **(E)**. Error bars represent s.d. of technical replicates (one representative of 2 experiments). DMSO and ARS-1620 data are shared between (C) and (E) as the combination experiments displayed were done side-by-side with the same ARS-1620 and DMSO controls. (**F)**, Annexin V-FITC and propidium iodide-based flow cytometry analysis of cell death in H358 cells treated with DMSO/KRASi (1 μM) together with GDC0941 (2 μM), erlotinib (1 μM) or 17-AAG (50 nM) (average of 2 or 3 experiments as indicated). Significance determined with t-test. **p<0.01, ***p<0.001.

### Identifying mechanisms of adaptation to KRASi in 2D vs. 3D growth conditions

Use of two-dimensional (2D) cultures to model adaptive responses to therapy can be a useful initial approach; however, three-dimensional (3D) models may better recapitulate in vivo physiology and drug sensitivity in some situations (Breslin and O’Driscoll, 2016; Imamura et al., 2015; Jacobi et al., 2017). Indeed, a subset of KRAS^G12C^-driven NSCLC cellular models demonstrate a differential sensitivity to KRASi in 2D versus 3D growth conditions and 3D growth conditions are more reflective of in vivo KRASi sensitivity (Fujita-Sato et al., 2015; Janes et al., 2018; Patricelli et al., 2016). To test the difference in drug response in 2D versus 3D, we chose two NSCLC cell lines (HCC44 and H2030) and confirmed at least 20-fold increased sensitivity to KRASi in 3D compared to 2D growth conditions (Janes et al., 2018) (Figure 6A, Figure S6A). Interestingly, basal pERK, pAKT and KRAS levels were all decreased in 3D versus 2D growth conditions (Figure 6B). KRASi induced further prominent decreases in pERK and pAKT, especially in the 3D condition (Figure 6B) (Janes et al., 2018; Riedi et al., 2017).

**Figure 6.**
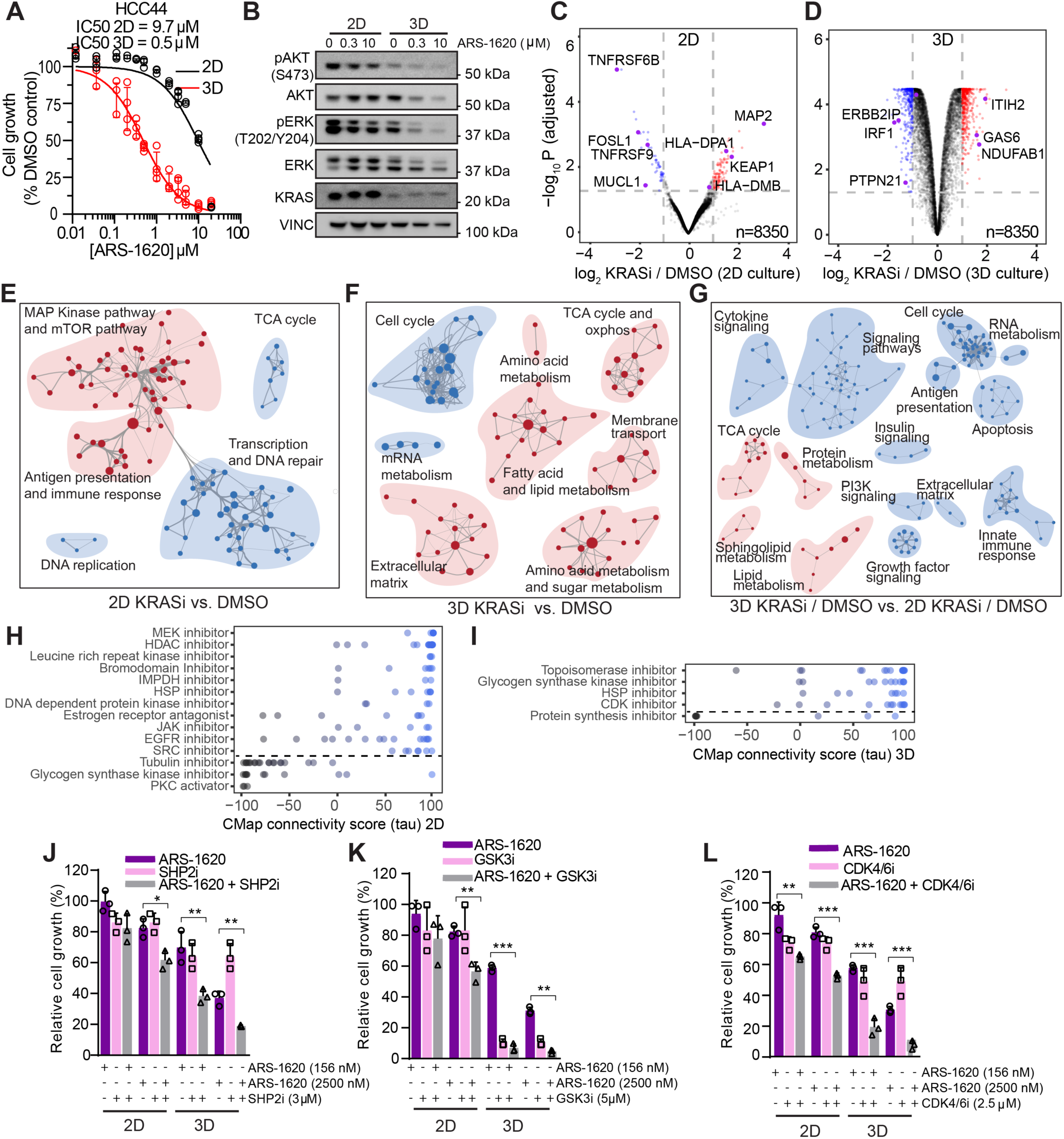
Quantitative proteomics of KRASi in 2D versus 3D growth conditions. **(A)** Cell proliferation dose-response curves for HCC44 cells treated with ARS-1620 (left: 2D, right: 3D). Error bars represent s.d. of 4 technical replicates (representative of 2 experiments). **(B)** Immunoblotting of lysates from HCC44 cells treated with KRASi (72 h) in 2D and 3D. **(C-D)** Volcano plots illustrate statistically significant protein abundance differences in HCC44 cells treated with KRASi in 2D vs. 3D for 72 h as in Fig. 1E. **(E-G)** Enrichment map of gene set enrichment analysis (GSEA) of KRASi-treated HCC44 cells versus DMSO in 2D **(E)** and 3D **(F)** or 3D compared to 2D **(G)** as in Fig. 1H. **(H and I)** Connectivity map analysis for 2D **(H)** and 3D **(I)** KRASi HCC44 proteomes as in Fig. 4A. **(J-L)** Relative cell growth for HCC44 cells treated with ARS-1620 vs. DMSO with or without SHP2i **(J)**, GSK3i **(K)** and CDK4/6I **(L)** in 2D vs. 3D. Data are from three independent replicates from a representative experiment of 3. The data used for SHP2i, GSK3i and CDK4/6i alone are the same within each 2D and 3D grouping and the bars are replicated for ease of comparison at the two different ARS-1620 dose levels. ARS-1620 alone data are the same between **(K)** and **(L)** as the GSK3i and CDK4/6i experiments were done side-by-side with the same ARS-1620 alone control. Significance determined with t-test vs. ARS-1620. *p<0.05, **p<0.01, ***p<0.001.

To identify differences in the baseline 2D versus 3D proteome and response to KRASi, we performed proteomic analyses of HCC44 cells treated with KRASi for 72 h (2D: 10 µM, 3D: 0.5 µM). Despite using growth condition-specific doses, KRASi in 3D growth conditions induced a larger number of significant proteomic alterations compared to 2D (Figure 6C-D, Table S7). GSEA of KRAS-inhibited 2D-HCC44 proteomes demonstrated an upregulation in the MAPK/mTOR pathway and antigen presentation (El-Jawhari et al., 2014) (Figure 6E) and downregulation of DNA replication/transcription/repair and TCA cycle (Figure 6E) all consistent with MiaPaCa-2 and H358 2D KRASi proteomic data. KRASi-treated 3D-HCC44 proteomes exhibited downregulation in cell cycle, a consistent response throughout all our datasets. In contrast to 2D data, KRASi in 3D-HCC44 led to upregulation in lipid and amino acid metabolism, TCA cycle and extracellular matrix pathways (Figure 6F).

To understand the basal differences in 2D versus 3D growth conditions and how this may have impacted response to KRASi (Figure S6B), we analyzed differential pathways by GSEA in basal 3D versus 2D proteomes (absent KRASi) (Figure S6C). In comparison to 2D, 3D growth conditions were associated with upregulation of antigen presentation, innate immune response, signaling pathways, carbohydrate metabolism and extracellular matrix, the latter two pathways likely upregulated as a result of increased HIF-1α stability in 3D growth conditions (Gagliano et al., 2016; Longati et al., 2013). Proteins associated with transcription and RNA metabolism pathways were downregulated in 3D versus 2D conditions. To identify KRASi responses independent of basal differences in 2D versus 3D conditions, we performed a comparative GSEA of 3D KRASi / 3D DMSO vs. the corresponding 2D conditions (Figure 6G). In response to KRASi, lipid/sphingolipid metabolism, TCA cycle and protein metabolism were differentially upregulated in 3D conditions while immune response/antigen presentation, growth factor/insulin signaling, cell cycle and RNA metabolism were downregulated. Interestingly, PI3K signaling was differently downregulated in 3D conditions, confirming the western blot data and previous studies with MAPKi showing a deeper inhibition of these pathways in 3D versus 2D growth conditions (Riedi et al., 2017). Overall, our data suggest that the mechanisms of adaptation are different in 2D versus 3D growth conditions. Given the high correlation observed between 3D culture conditions and in vivo susceptibility to drugs, identification of mechanisms of adaptation in 3D might have a greater therapeutic relevance.

Next, we performed CMap analysis to identify drugs that correlated with the proteomic profile in 2D and 3D growth conditions. HCC44-2D analysis revealed similar compounds as in MiaPaCa-2 and H358 KRASi treated cells, including MEKi, IMPDHi, HSP90i, EGFRi, and HDACi (Figure 6H). In the HCC44-3D analysis, HSP90i were enriched, in common with 2D analyses. Specific to the 3D KRASi, we identified a positive correlation with CDKi and topoisomerase inhibitors (Figure 6I). Interestingly, glycogen synthase kinase inhibitors (Kazi et al., 2018) positively correlated with 3D-KRASi (Figure 6I) but were negatively correlated with the 2D-KRASi (Figure 6H), suggesting these inhibitors might have opposite effects in 3D versus 2D growth conditions. We next performed CMap analysis of DMSO treated cells in 3D vs 2D conditions as well as 3D KRASi to DMSO vs. 2D KRASi to DMSO (Figure S6D), which reinforced the positive correlation with GSK3i response in 3D-KRASi.

Finally, we tested the efficacy of select KRASi combinations in 2D and 3D growth conditions (SHP2i, GSK3i, CDK4/6i). We tested SHP2i given reactivation of MAPK signaling as determined in our GSEA and positive results of combination SHP2i with MEKi (Wong et al., 2018). Interestingly, SHP2i alone was effective in 3D compared to 2D growth conditions across multiple cell lines (Figure 6J, Figure S6E) and the combination with KRASi induced a further response in 3D conditions. GSK3i (CHIR-99021) was also more effective in 3D versus 2D conditions in HCC44 cells but not in H2030 highlighting the importance of testing candidate combinations in multiple cellular models (Figure 6K, Figure S6F). Finally, as predicted by GSEA and CMap analysis, KRASi combined with CDK4/6 inhibition had more potent effects on cell growth inhibition in 3D compared to 2D across multiple cell lines (Figure 6L, Figure S6G). These results suggest that quantitative proteomics can identify differential mechanisms of adaptation to KRASi in 2D versus 3D growth conditions.

## DISCUSSION

After years of considering oncogenic KRAS an undruggable target, new small-molecule inhibitors targeting KRAS^G12C^ are showing potent target engagement and efficacy in early phase clinical trials. Despite these exciting preliminary results, driver oncoprotein targeted drug efficacy is usually limited by tumor heterogeneity and by compensatory adaptive mechanisms. Here, we modeled the effects of acute and long-term KRAS^G12C^ inhibition to investigate non-mutational mechanisms of adaptation in KRAS^G12C^ mutant PDAC and NSCLC cellular models. We identified shared acute and long-term KRASi proteomic signatures associated with persistent cell survival and designed rational combinatorial strategies to induce cytotoxicity thereby laying the foundation for in vivo pre-clinical testing. Our results suggest that deep quantitative temporal proteomics combined with bioinformatic analyses is one useful method to identify new combination therapies to circumvent or prevent development of resistance to targeted therapies.

The shared overarching acute and long-term response to KRASi as determined by our proteomic and phosphoproteomic analyses and confirmed by decades of prior work consists of first, an acute cytostatic response with downregulation of cell cycle, transcription, translation, and pro-growth signaling pathways concurrent with upregulation of compensatory stress response and metabolic pathways (Fig. 7). In cellular models that are able to re-establish proliferation despite ongoing KRASi, our long-term KRASi proteomic signatures revealed upregulation of cell cycle pathways, transcription, and DNA repair relative to acute KRASi. Furthermore, the long-term KRASi proteomic signature was marked by altered cellular metabolism (upregulation of mitochondrial respiration) with a prominent decrease in mTOR signaling with coincident upregulation in lysosomal metabolism, which likely correlates with a compensatory activation of autophagy also described in ERK-treated PDAC (Bryant et al., 2019). The fact that oncogenic KRAS remodels cellular metabolism is well known (Bryant et al., 2014) and our data is in agreement with studies of genetic ablation of KRAS in PDAC which leads to a shift towards mitochondrial metabolism and autophagy (Viale et al., 2014). Long-term KRASi also altered extracellular matrix pathways likely related to enhanced adherence and dependence of adhesion in KRAS-ablated cells (Chen et al., 2018) (Tape et al., 2016).

**Figure 7.**
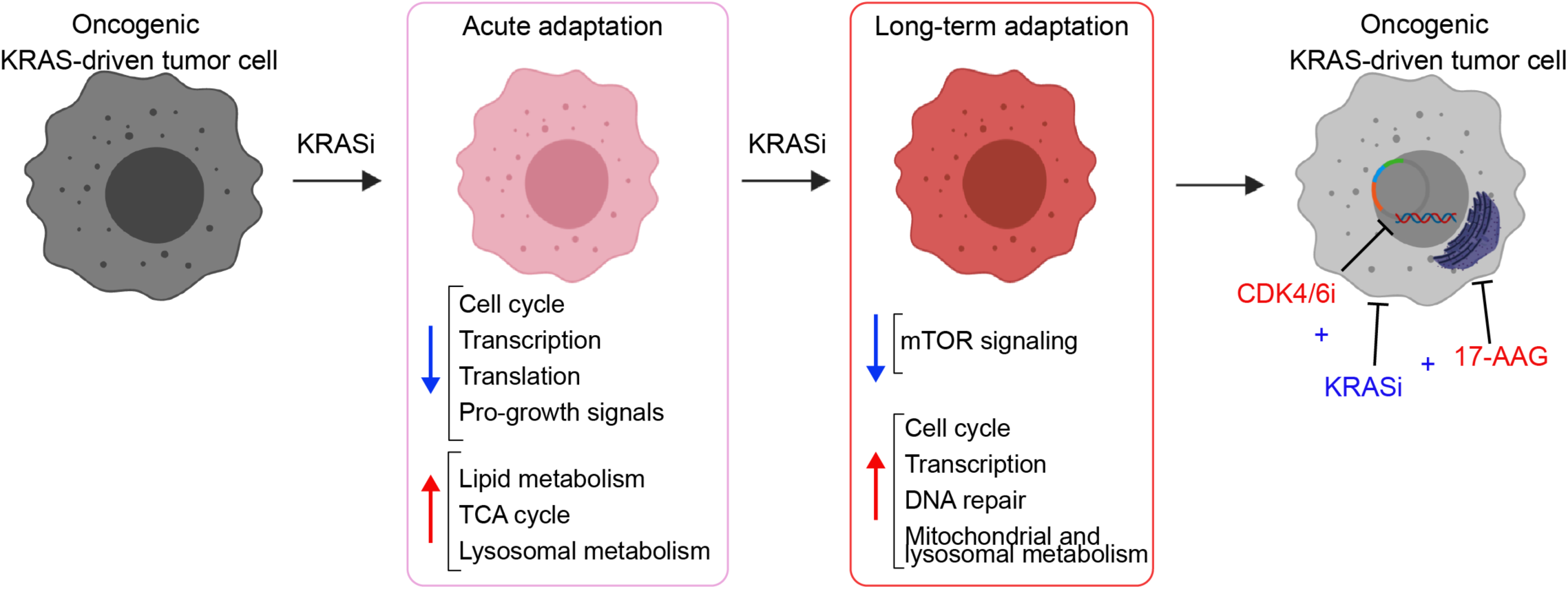
KRASi acute and long-term adaptation. Oncogenic KRAS-driven cells acutely treated with KRASi or after developing long-term adaptation to KRASi were analyzed using quantitative proteomics. Bioinformatic analyses determined significant up- and downregulated pathways at each timepoint. Acute adaptation consists of a cytostatic response with downregulated cell cycle, transcription, translation and pro-growth signals and concomitant alteration of metabolic pathways with upregulation of lipid metabolism, TCA cycle and lysosomal processes. Long-term adaptation consists of re-established proliferation coincident with increased cell cycle, transcription, DNA repair and mitochondrial metabolism together with consistent downregulation of mTOR signaling and increased lysosomal processes. Subsequent bioinformatic analyses nominated high-priority KRASi combinatorial drug strategies. KRASi combinations with HSP90i (17-AAG) and cell cycle inhibitors (CDK4/6i) efficiently blocked growth in 2D and 3D culture models.

The large number of pathway alterations in response to KRASi is unsurprising given the dependence on oncogenic KRAS to support tumor proliferation and survival. Interestingly, in comparing the proteomic signature of KRASi with a distinct treatment (GLSi) that likewise causes an acute cytostatic response with eventual re-establishment of proliferation, a large number of pathways were found in common at both acute and long-term timepoints. For instance, KRASi and GLSi (Biancur et al., 2017) both lead to an induction of a differential metabolic rewiring consisting in activation of mitochondrial respiration, lipid metabolism and lysosomal function. This suggests that despite vastly different targets and mechanisms of action, many aspects of the proteomic response to treatments that induce cytostasis are shared. Indeed adaptation to pharmacologic KRASi correlated with a long-term reactivation of cell cycle and DNA repair as well as MYC levels, all pathways also activated in gemcitabine-resistant (Seth et al., 2019), ERKi-resistant, and genetic KRASi-resistant cells (Santana-Codina et al., 2018; Vaseva et al., 2018). This finding is further highlighted by results from our CMap analysis that identify a large number of shared signatures with KRASi at acute time points. While several of these shared signatures are due to inhibition of downstream aspects of the KRAS pathway (MEKi), the unrelated drug classes and sheer number of shared responses suggests that the shared CMap signature is at least in part due to a generalized cytostatic response. One hypothesis based on these results is that co-targeting pathways that share an overall compensatory response signature associated with cytostasis may overwhelm these shared or parallel compensatory pathways and tip the balance from cytostasis to cytotoxicity. To evaluate this hypothesis, we tested a number of KRASi combinations and demonstrated potentially useful therapeutic combinations that prevented re-establishment of proliferation.

Successful KRASi combinations across multiple cellular models included combination with PI3Ki, HSP90i, CDK4/6i, and SHP2i. Co-targeting KRAS^G12C^ with PI3Ki reduced proliferation in long-term growth assays and induced cytotoxicity, in agreement with multiple previous studies (Alagesan et al., 2015; Engelman et al., 2008; Misale et al., 2018; Muzumdar et al., 2017) and validating the efficacy of our approach. Given the toxicity and poor response of patients with combinatorial targeting of KRAS/MEK and PI3K pathways (Janne et al., 2017; Pettazzoni et al., 2015; Will et al., 2014), we also evaluated multiple additional combinations. KRASi in combination with HSP90i (17-AAG) also induced cytotoxicity in multiple KRAS^G12C^-driven cellular models. This is consistent with increased sensitivity to ER stress induction in KRAS-resistant PDAC cells (Genovese et al., 2017), likely due to efficient block of the reported pERK surge after 17-AAG treatment (Zhang et al., 2010). This combination is particularly promising for future pre-clinical in vivo testing as it targets a completely distinct pathway from canonical downstream KRAS signaling. Finally, CDK4/6i and SHP2i worked in combination with KRASi in multiple 3D cultures supporting recent studies that tested its potential in combination with MEKi (Haines et al., 2018; Mainardi et al., 2018; Ruess et al., 2018). Indeed, in vivo pre-clinical evaluation of CDK4/6i in combination with KRASi showed promising results (Lou et al., 2019). Overall, our approach was able to predict useful combinations with KRASi and drugs targeting multiple pathways that will benefit from further evaluation in pre-clinical models.

Given the robust nature of our KRASi proteomics data that correlates in full with decades of research, the unique temporal proteomics aspect, and 2D versus 3D growth condition proteomic comparisons, we have generated an interactive KRASi proteome database accessible via an easy to use website (https://manciaslab.shinyapps.io/KRASi/). This a resource for the oncogenic KRAS biology community that allows for systematic search by gene name, cell line or time of treatment within our KRASi proteomics datasets (Figure S7). With KRAS^G12C^ inhibitors now being tested in clinical trials with promising results and more being developed targeting different mutant forms (Kessler et al., 2019), anticipating the potential mechanisms of resistance and developing useful combination therapies is critical. Our study uncovers a large number of compensatory proteomic responses to KRASi that at first suggest a daunting amount of collateral survival pathways employed by oncogenic KRAS dependent pathways. However, with combination targeting of these pathways, this suggests that this large number of compensatory pathway activations actually reveal a large number of liabilities for combination targeting. Although more studies are required to understand the potential translatability of our findings using in vivo pre-clinical models, our proteomics platform has the capacity to define the proteomic response to any drug and facilitate the design of rational combinatorial treatment that might ultimately represent a new therapeutic approach for patients.

## ACKNOWLEDGEMENTS

We gratefully acknowledge Dr. Steven Gygi for use of CORE for mass spectrometry data analysis software. This research has been supported by grants from Burroughs Wellcome Fund Career Award for Medical Scientists, Brigham and Women’s Hospital MFCD Award, Sidney Kimmel Foundation Kimmel Scholar Program, Dana-Farber Cancer Institute Claudia Adams Barr Program for Innovative Cancer Research Award and the Hale Family Center for Pancreatic Cancer Research to J.D.M. A.S.C is supported by a Canadian Institute of Health Research (CIHR) post-doctoral fellowship. W.F. and B.M. are supported by the First TEAM of the Foundation for Polish Science and research funds of the Medical University of Lodz (503/1-090-03/503-11-001-19-00).

## AUTHOR CONTRIBUTIONS

J.D.M., A.S.C., and N.S.C. conceived the study and designed experiments. A.S.C., N.S.C., and A.G. performed all experiments. TMT-based quantitative proteomics experiments were also performed by M.K., M.P.J., and S.G. Q.Y., B.M., M.S., and W.F. performed bioinformatics analysis of proteomic data. J.D.M, A.S.C. and N.S.C. analyzed the data and wrote the manuscript. All authors edited the manuscript.

