## Supplementary Information for "Defining and targeting adaptations to oncogenic KRAS^G12C^ inhibition using quantitative temporal proteomics"

#### SUPPLEMENTARY METHODS

**Cell Culture.** Mia-PaCa2, H358, and H2030 cells were purchased from the American Type Culture Collection (ATCC). HCC44 was purchased from DSMZ. Cell lines were carefully maintained in a centralized cell bank, and routinely inspected for mycoplasma contamination using PCR. All cell lines were maintained in a humidified incubator at 37C with 5% $CO_2$ and grown in DMEM (Invitrogen 11965) or RPMI 1640 (Invitrogen 11875), supplemented with 10% Fetal Bovine Serum (FBS) and 1% Penicillin/Streptomycin. Cells were grown on standard tissue culture plates for 2D culture or on ultra-low attachment plates for 3D culture.

**Cell Proliferation Assay.** Cells were plated at 3000-5000 cells per well in 24-well plates. The next day, cells were treated with FRF-01-157 or ARS-1620 (dose as indicated). Of note, similar results were obtained whether the media was refreshed or not refreshed with addition of new compound. Cells were fixed at indicated time points, with 10% formalin followed by staining with 0.1% crystal violet. Dye was extracted with 10% acetic acid, and relative proliferation was determined by measuring OD at 595nm. Data was normalized to day 0 readings.

**IC50 Assay.** Cells were plated in 96-well plates at 2000 cells per well and treated with the indicated compounds at serial dilutions the next day. Cell viability was measured after 72 hours, using Cell-Titer Glo (Promega, G7570) assay. IC50 value was calculated using non-linear regression analysis (log(inhibitor) vs. normalized response) using Graphpad Prism.

**Cell death assay.** Flow cytometry analysis of apoptosis: Cells were plated and treated for 72h with the indicated compounds. Cells were stained with Annexin-FITC and propidium iodide (PI) for 15 min (BD Biosciences 556547) using the manufacturer's protocol. Cells were placed in ice and analyzed using a Beckman Coulter Cytoflex.

**Western Blot Analysis.** Cells were collected in ice cold PBS at a confluency of 80-90%. Cells were lysed in RIPA buffer (50 mM TrisHCl pH 7.4, 150 mM NaCl, 2 mM EDTA), containing 1% NP-40, 0.1% SDS, protease and phosphatase inhibitors (Life Technologies, 78847) and 1 mM DTT. Lysates were centrifuged at 14000 X g for 15 minutes at 4°C. Supernatants of the cell lysates were collected into separate Eppendorf tubes. Protein concentration determination was performed using the bicinchoninic acid (BCA) assay (Thermo Fisher, 23250). 30-50 μg of protein was resolved on 4-12% SDS-PAGE gels (Thermo Fisher, NP0322) and transferred to polyvinylidene difluoride (PVDF) membranes (Bio-Rad). The membranes were first blocked in Tris-buffered saline with 0.1% Tween 20 (TBS-T) containing 5% milk for an hour. The membranes were incubated with specific primary antibodies overnight at 4°C. Membranes were washed three times with TBS-T, followed by incubation with peroxidase-conjugated secondary antibody (1:10000) for 1 h. Membranes were developed using the Enhanced Chemiluminescence (ECL) Detection System (Thermo Fisher, 32209). The antibodies used are as follows: p-AKT (Ser473, Cell Signaling #4060S, 1:1000), AKT (Cell Signaling #9272, 1:1000), pERK1/2 (Thr202/Tyr204, Cell Signaling #4376, 1:1000), ERK1/2 (Cell Signaling #9102, 1:1000), KRAS (Santa Cruz F234 sc-30, 1:100), Vinculin (EMD Millipore v284, 1:5000), Anti-rabbit IgG (H1L) HRP conjugate (Thermo 31460, 1:3000); anti-mouse IgG HRP conjugate (Promega W402B, 1:5000).

**Metabolomics.** Steady state metabolomics experiments were performed as previously described (Son et al., 2013; Sousa et al., 2016). Briefly, MiaPaCa-2 cells were plated and media was refreshed the next day with drug/DMSO. Metabolite collection was performed after 24 h, 72 h or

8 weeks treatment. Experiments were performed in 25 mM glucose/4 mM glutamine. Metabolite fractions were normalized to cell number obtained in a parallel 6 cm plate.

**Quantitative Proteomics.** Cells were plated and compounds were refreshed for long-term experiments. Cells were harvested in ice-cold PBS. Quantitative mass spectrometry-based proteomics were performed as previously described (Biancur et al., 2017; Paulo et al., 2017). TMT isobaric reagents were from Thermo Scientific (909406, Life Technologies). Water and organic solvents were from J.T. Baker (Center Valley, PA, USA). Cell lysis was carried out in 2% SDS buffer with 25 mM NaCl, 20 mM HEPES pH 8.8, 5 mM dithiothreitol (DTT, to reduce disulfide bonds), 200 µM sodium orthovanadate (New England Biolabs, P0758S), and 1 x Halt™ Protease and Phosphatase Inhibitor Cocktail (Life Technologies, 78847) with zirconium oxide beads (Next Advance / MidSci ZROB05) using a vortexer at maximum speed for 5 min (Tissue Lyser LT, Qiagen). The homogenized cell lysate was placed at 60°C for 30 min, followed by cooling at room temperature for 10 min. Proteins were alkylated with 14 mM iodoacetamide (I1149 Sigma) at room temperature for 30 min in the dark. Excess iodoacetamide was quenched with 15 mM dithiothreitol at room temperature for 15 min in the dark.

Chloroform-methanol precipitation of proteins from cells was performed prior to protease digestion. To summarize, three parts neat methanol was added to each sample and vortexed, one part chloroform was added to the sample and vortexed, and 2.5 parts water was added to the sample and vortexed. The sample was centrifuged at 4000 x g for 10 min at room temperature and subsequently washed twice with 100% methanol. Samples were resuspended in 200 mM HEPES pH 8.5 for digestion. Protein concentrations were determined using the bicinchoninic acid (BCA) assay.

For each sample, 100 µg of protein was digested overnight with 1:100 protease-to-protein ratio Lys-C protease (129-02541, Wako Chemicals USA, Inc.) followed by trypsin (V5117, Promega) at a 1:100 protease-to-protein ratio, all at 37°C. Digested peptides were separated from contaminants using 50 mg silica bonded phase cartridges (WAT054960, Waters) followed by lyophilization using a speedvac concentrator overnight. Approximately 50 µg of peptides from each sample were labelled with TMT reagent (100 µg) for 1 h, followed by quenching the reaction with hydroxylamine to a final concentration of 0.3% v/v. The TMT-labeled samples were combined at a 1:1:1:1:1:1:1:1:1:1:1 ratio. The combined sample was acidified, vacuum centrifuged to near dryness and subjected to C18 SPE (Sep-Pak, Waters). Samples were separated using basic pH reversed-phase HPLC and then pooled into 12 fractions. Data were obtained using an Orbitrap Fusion Lumos mass spectrometer (Thermo Fisher Scientific, San Jose, CA, USA) coupled with a Proxeon EASY-nLC 1200 LC pump (Thermo Fisher Scientific). Peptides were separated on a 75 µm inner diameter microcapillary column packed with 35 cm of Accucore C18 resin (2.6 µm, 100 Å, Thermo Fisher Scientific).

For phospho-peptide analyses, sample preparation was performed using 1 mg per sample and was as described above up to protein digestion with LysC and trypsin. After protein digestion, samples were desalted using 200 mg silica bound phase cartridges followed by lyophilization using a speedvac concentrator. Next, each sample was enriched for phospho-peptides using the Pierce High-Select FE-NTA phosphopeptide Enrichment Kit following the manufactures instructions. After phospho-peptide elution, each sample was lyophilized using a speedvac concentrator. Phospho-peptides were resuspended in 200 mM EPPS buffer and a BCA assay was performed. TMT-labeling of phospho-peptides was performed as described earlier. Samples were mixed in 1:1:1:1:1:1:1:1:1:1:1 ratio and the combined sample was vacuum centrifuged to near dryness and subjected to C18 SPE. The combined phospho-peptide sample was further fractionated using basic pH reversed-phased as described earlier into 24 pooled fractions and data were acquired using an Orbitrap Fusion Lumos mass spectrometer. Peptides were separated using a 3 hour gradient of 6-27% acetonitrile in 0.125% formic acid with a flow rate of 400 nL/min. Each analysis used an $MS^3$-based TMT method as described previously (McAlister et al., 2014). The data were acquired using a mass range of m/z 350-1350, resolution 120,000, AGC target 1 X $10^6$, maximum injection time 100 ms, dynamic exclusion of 120 s for the peptide measurements in the Orbitrap. Data dependent $MS^2$ spectra were acquired in the ion trap with a normalized collision energy (NCE) set at 35%, AGC target set to 1.8 X $10^4$ and a maximum injection time of 120 ms. $MS^3$ scans were acquired in the Orbitrap with a HCD collision energy set to 55%, AGC target set to 1.5 X $10^5$, maximum injection time of 150 ms, resolution at 50,000 and with a maximum synchronous precursor selection (SPS) precursors set to 10. Phospho-peptides were analyzed using a 2 h gradient of 1-22% acetonitrile in 0.125% formic acid with a flow rate of 400 nL/min and were acquired using the same parameters as described above with the addition of MultiStage Activation (MSA). Mass spectra were processed using a Sequest-based-in-house software pipeline as described previously(Paulo et al., 2015). Protein quantitation values were exported for further analysis. Each reporter ion channel was summed across all quantified proteins and normalized assuming equal protein loading in all 9-11 samples.

**Bioinformatic Analysis.** Quantitative proteomics data was further processed with LIMMA package (3.40.2) (Ritchie et al., 2015) in R platform. Pairwise comparisons were performed among experimental conditions. A standard linear model fitting and an empirical Bayes procedures were performed to correct the distribution. A moderated t-statistic using a simple Bayesian model was applied as the fundamental statistic, followed by multiple test correction through the Benjamini-

Hochberg method, which is referred to as false discovery rate (FDR). All these procedures are performed via limma with default configuration. For each comparison, a volcano plot was used to visualize significantly different proteins. Gene set enrichment analysis (GSEA) of each comparison was performed with Broad GSEA software (3.0) (Subramanian et al., 2005 , PNAS) using the collection containing all canonical pathways (c2.cp.v6.2.symbols.gmt) in MSigDB for pathway annotation. GSEA results were further visualized with EnrichmentMap (3.2.1) (Merico et al., 2010) in Cytoscape (3.7.2) (Shannon et al., 2003), and enriched pathway clusters were manually curated. Gene Ontology (GO) enrichment analyses were performed by using two unranked lists of genes, target list and background lists as previously described (Eden et al., 2009). In principal component analysis (PCA), the fold changes of proteins in KRASi-treated samples vs. DMSO controls were calculated across different time points and were used as input datasets for the analysis. PCA results were visualized in a two-dimensional coordinate space according to two major principal components.

To retrieve compound inhibitors with similar expression signature, the top 150 up- and down-regulated proteins selected and queried against the L1000 gene expression database in Connectivity Map (CMap) 2.0 (Subramanian et al., 2017). The compound inhibitor classes with connectivity score greater than 90 or less than -90 were identified.

Significantly increased and decreased proteins in each comparison were queried against DrugBank (Version 5.1.4)(Wishart et al., 2018), and proteins with experimental drugs or FDA-approved drugs were annotated.

For overlap analysis of the KRASi proteomic data with the RNA-seq and GSEA analyses published by Janes et al. (Janes et al., 2018), enrichment of the described gene sets was analysed in the KRASi proteome data with GSEA v3.0  (Subramanian et al., 2005 , PNAS). Analysis of signatures was performed using a separate GSEA run with 1,000 permutations (gene set) with default ranking parameters.

To further understand the shared KRASi proteomic signature between MiaPaCa-2 and H358 KRASi proteomes, a meta-analysis of pathway and protein-protein interaction (PPI) enrichment was performed using Metascape (Zhou et al., 2019). The top 300 up- or down-regulated proteins were selected with FDR smaller than 0.05 by LIMMA algorithm. Proteins identified across cell lines were used as background input for pathway enrichment analysis. Protein-protein interaction networks were queried against BioGRID, InWeb_IM and OmniPath databases via the Metascape platform. Function-related sub-networks were clustered by Molecular Complex Detection (MCODE) algorithm (Bader and Hogue, 2003), and manually annotated. Data processing and statistical analysis were performed on R (3.5) platform.

**Chemicals**. ARS-1620 (Chemgood, C-1454), Compound 4 (Zeng et al., 2017), 17-AAG (Cayman chemicals 75747-14-7), GDC-0941 (Selleckchem, S1065), PF 2341066 (crizotinib, Cayman chemicals, 877399-52-5), Erlotinib hydrochloride (Cayman chemicals, 183321-74-6), CINK4 (CDK4/6 inhibitor, Cayman chemicals, 359886-84-3), SHP099 hydrochloride (SHP2i, Selleckchem, S8278), CHIR-99021 (GSK3α/βi, Selleckchem S1263).

**Statistics.** All the data except the proteomics (analysis described above) was analyzed using GraphPad PRISM software. No statistical methods were used to predetermine sample size. For comparison between two groups, Student's t-test (unpaired, two-tailed) was performed for all experiments. Groups were considered different when $p<0.05$.

#### SUPPLEMENTARY FIGURE LEGENDS

**Figure S1. Quantitative temporal proteomics to determine mechanisms of adaptation to KRASi in PDAC cells. (A)** Volcano plot illustrates statistically significant protein abundance differences in MiaPaCa-2 cells treated with KRASi for 4 h. Volcano plots display the $-\log_{10}$ (P value) versus the $\log_2$ of the relative protein abundance of mean KRASi treatment to mean control (DMSO). Red circles represent significantly (p value/FDR $\leq$ 0.05) upregulated proteins (FC $\geqslant$ 2) while blue circles represent the significantly downregulated proteins (FC $\leq$ 0.5). Data are from three independent plates from a representative experiment of 2. **(B and C)** Enrichment map of gene set enrichment analysis (GSEA) of KRASi-proteomes from MiaPaCa-2 cells at 24 h (IC90 dose) **(B)** and 8 weeks **(C).** FDR<0.01, Jaccard coefficient>0.25, node size is related to the number of components identified within a gene set and the width of the line is proportional to the overlap between related gene sets. GSEA terms associated with upregulated (red) and downregulated (blue) proteins are colored accordingly and grouped into nodes with associated terms**. (D-H)** Temporal GSEA of MiaPaCa-2 cells treated with KRASi for 1 h, 4 h, 24 h, 72 h, 7 d and 8 weeks showing top downregulated and upregulated pathways at 24 h **(D, F)** and 7 days **(E, G)** and lysosome upregulation and mTOR downregulation **(H)**.

**Figure S2. Metabolic adaptations to acute and long-term KRASi.** For all metabolomics experiments in this figure, cells were treated with Compound 4 (5 μM) for 24 h, 72 h or ARS-1620 (2 μM) for 8 weeks in media containing glucose (25 mM) and glutamine (4 mM). Pathways were decreased at 24 h but increased at 72 h and maintained at 8 weeks. Values are represented as fold change versus DMSO-treated cells, significant metabolites from each pathway are represented.

Error bars represent s.d. of n=3 technical replicates from independently prepared samples from individual wells. Significance determined for cells treated with KRASi vs. vehicle. **(A)** Fold change of glycolytic intermediates after KRASi vs. DMSO. G6P, glucose 6-phosphate; G1P. glucose 1-phosphate; F6P, fructose 6-phosphate; FBP, fructose 1,6-bisphosphate; Ga3P, glyceraldehyde 3-phosphate; 3PG, 3-phosphoglycerate; PEP, phosphoenolpyruvate; Lac, lactate. **(B)** Fold change of metabolites in the hexosamine biosynthesis pathway (HBP). Glc6P, Glucosamine-6P, GlcNAc-1P, N-Acetyl-glucosamine-1-phosphate; UDP, uridine diphosphate; UDP-GlcNAc, UDP-N-Acetyl-glucosamine, UDP-G, UDP-glucose. **(C)** Fold change of metabolites in the non-oxidative Pentose Phosphate Pathway (PPP). DHAP, dihydroxyacetone phosphate; S7P, seduheptulose-7P, SBP, seduheptulose-1,7-bisP; OBP, Octulose-1,8-phosphate. **(D)** Fold change of metabolites in the pyrimidine and purine pathways after KRASi treatment. UMP, uridine monophosphate; dCTP, deoxy cytidine triphosphate; dTTP, deoxythymidine triphosphate; IMP, inosine monophosphate; dAMP, deoxyadenosine monophosphate; CMP, cytidine monophosphate; IDP; inosine diphosphate; dGTP, deoxyguanosine triphosphate; ATP, adenosine triphosphate. Significance determined with t-test. *$p<0.05$, **$p<0.01$, ***$p<0.001$.

**Figure S3: Phosphoproteomic analysis of signaling changes in KRASi treated cells. (A-C)** Volcano plots illustrate statistically significant protein abundance differences in MiaPaCa-2 cells treated with KRASi for 1 h, 4 h and 24 h. Volcano plots display the $-\log_{10}$ (P value) versus the $\log_2$ of the relative phospho-peptide abundance of mean KRASi treatment to mean control (DMSO). Red ($\log_{10}FC \geq 1$) and blue ($\log_{10}FC \leq -1$) dots indicate significant (p value $\leq 0.05$) up- and downregulated phospho-peptides. Data from each time point are from three independent plates from one experiment. **(D)** Heatmap of DMSO and KRASi cells for 24 h. Rows represent different phospho-peptides and color represents directionality of expression. Intensity correlates with Z-score (deviation from each phospho-peptide mean across all samples) for each sample. **(E-F)** Gene Ontology analysis of the top down- **(E)** and upregulated **(F)** phospho-peptides at 24 h of KRASi classified by p-value. Vertical lines indicate adjusted p-value=1.3. **(G)** The top up- and downregulated phospho-peptides were selected at 24 h and analyzed at different time points. For all proteins, multiple phospho-peptides were identified. Peptide selection was performed based on top fold change and relevance to cancer and therapeutic resistance based on literature search (Phosphosite). Phospho-peptides were normalized to whole proteome to show net change in phosphorylation independent of changes in protein expression.

**Figure S4: KRASi proteomics correlation with KRASi genetic studies and CB-839 proteomics. (A and B)** Correlation plot of MiaPaCa-2 KRASi proteomic datasets at 24 h **(A)** and 7 d **(B)** with a CRISPRi genome screen (Lou et al., 2019) showing poor correlations. **(C)** Heatmap of top 150 proteins from H358 and MiaPaCa-2 KRASi proteomics and mRNAs from H358 RNA-seq data (Janes et al., 2018). **(D)** Normalized expression scores (NES) of KRASi proteomics and RNA-Seq gene set enrichment analysis (GSEA hallmarks (MSigDb database c1.v6.2 Hallmark). **(E)** Venn diagram of GSEA pathways identified in MiaPaCa-2 KRASi (blue) and MPDAC-4 CB-839 (pink) treated cells at each time point shows overlap of dysregulated pathways. **(F-H)** Enrichment map of overlapped GSEA terms between KRASi-treated MiaPaCa-2 cells and CB-839-treated MPDAC4 at 24 h **(F)**, 72 h **(G)** and long-term **(H)** (all standardized to respective DMSO controls). For CB-839 GSEA analysis FDR<0.15, coefficient overlap (CO)>0.1 was used as previously published (Biancur et al., 2017) and for MiaPaCa-2 FDR<0.01 and Jaccard coefficient>0.25 was used. Node size is related to the number of components identified within a gene set and the width of the line is proportional to the overlap between related gene sets. GSEA terms associated with upregulated (red) and downregulated (blue) proteins in both MiaPaCa-2 and CB-839 are colored accordingly and grouped into nodes with associated terms. Isolated dots in right lower corner of Figure S4G belong to uncategorized GSEA terms.

**Figure S5: Targeting adaptation to KRASi in pancreatic and lung cancer. (A)** Cell proliferation dose-response curves for MiaPaCa-2 cells treated with 17-AAG, crizotinib and GDC0941. Error bars represent s.d. of 4 technical replicates (one representative of 2 experiments). **(B)** Immunoblot of lysates from MiaPaCa-2 cells treated with ARS-1620 for indicated times. **(C)** Cell proliferation dose-response curves for H358 cells treated with erlotinib and GDC0941. Error bars represent s.d. of 4 technical replicates (one representative of 2 experiments).

**Figure S6: Defining drug combinations with KRASi in 2D and 3D. (A)** Cell proliferation dose-response curves for H2030 cells treated with ARS-1620 in 2D and 3D growth conditions. Error bars represent s.d. of 4 technical replicates (one representative of 3 experiments). **(B)** Volcano plots illustrate statistically significant protein abundance differences in HCC44 cells treated with DMSO in 2D vs. 3D growth conditions for 72 h as in Fig. S1A. **(C)** Enrichment map of gene set enrichment analysis (GSEA) of DMSO-treated HCC44 cells in 3D vs. 2D as in Fig. S1B. **(D)** Connectivity map analysis for HCC44 cells in 3D vs. 2D with DMSO **(top)** or with KRASi **(bottom)**. Perturbagen classes with mean connectivity scores >90% or <-90% and FDR<0.05 are displayed. **(E-G)** Relative cell growth for H2030 cells treated with ARS-1620 vs. DMSO with or without SHP2i **(E)** and GSK3i **(F)** and CDK4/6i **(G)** in 2D vs. 3D. Data are from three independent replicates from a representative experiment of 2. The data used for SHP2i, GSK3i and CDK4/6i alone are the same within each 2D and 3D grouping and the bars are replicated for ease of comparison at the two different ARS-1620 dose levels . ARS-1620 alone data are the same between **(E-G)** as the combination experiments displayed were done side-by-side with the same ARS-1620 alone control. Significance determined with t-test vs. ARS-1620 **p<0.01, ***p<0.001.

**Figure S7: KRASi proteome website interface** https://manciaslab.shinyapps.io/KRASi/ The "KRASi proteome" is an interactive website that includes data from quantitative temporal KRASi proteomic datasets used in this analysis. This resource allows search by gene name, cell line or time of treatment within the KRASi proteomics datasets.

**Figure S8:** Uncropped immunoblots for all data elements shown in main figures.

**SUPPLEMENTARY TABLES:**

**Table S1: List of KRASi quantitative temporal proteomic experiments.**

**Table S2: Gene set enrichment analysis (GSEA) of MiaPaCa-2 KRASi quantitative temporal proteomic comparison at 24 h and 7 days.** Gene sets with a false discovery rate <0.01 and coefficient score>0.25 were entered into enrichment map analysis (Fig. 1H-J, S1C). From left to right as labeled, gene sets associated with downregulated proteins and gene sets associated with upregulated proteins.

**Table S3. Proteome and phosphoproteome of MiaPaCa-2 cells treated with KRASi for 1 h, 4 h, and 24 h.** TMT proteomic raw data and comparisons between DMSO- and KRASi-treated MiaPaCa-2 cells treated for 1 h, 4 h or 24 h. False discovery rates (FDRs) were calculated using the Benjamini-Hochberg procedure. Columns include: Uniprot protein identification number (proteinID), gene symbol (Gene Symbol), protein description/name (Description), number of peptides quantified per protein (peptides), the normalized summed signal-to-noise (sn) for each of the 10 channels. The fold-change comparisons of the KRASi treated to DMSO treated samples and Benjamini-Hochberg calculated FDRs can be found for each time point in separate tabs for proteome and phosphoproteome data.

**Table S4. Gene set enrichment analysis (GSEA) of H358 KRASi quantitative temporal proteomic comparison at 24 h and 7 days.** Gene sets with a false discovery rate <0.01 and coefficient score>0.25 were entered into enrichment map analysis (Fig. 2F-H). From left to right as labeled, gene sets associated with downregulated proteins and gene sets associated with upregulated proteins.

**Table S5. Connectivity scores from CMap analysis using top 150 up- and downregulated proteins.**

**Table S6. Identification of cell surface proteins and druggable targets with FDA-approved drugs.** Significantly increased/decreased proteins in each comparison were queried for identification of cell surface proteins and against DrugBank (Version 5.1.4 ) proteins to identify druggable targets with experimental drugs or FDA-approved drugs.

**Table S7. Gene set enrichment analysis (GSEA) of HCC44 KRASi quantitative temporal proteomics in 2D vs. 3D.** Gene sets with a false discovery rate <0.01 and coefficient score>0.25 were entered into enrichment map analysis (Fig. 6E-G, S6C). From left to right as labeled, gene sets associated with downregulated proteins and gene sets associated with upregulated proteins.

#### Figure S1

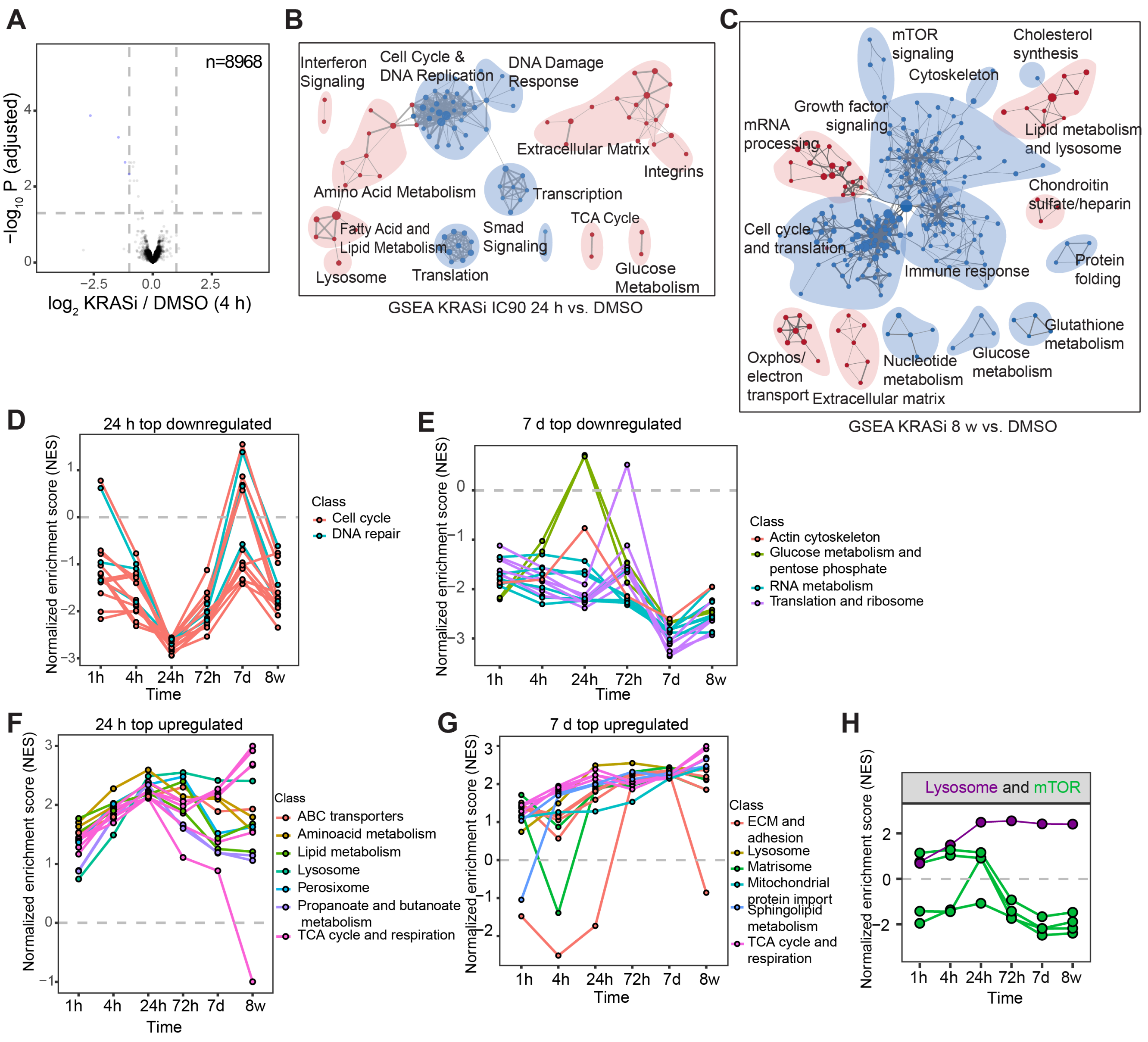

A
n=8968
−log10 P (adjusted)
log2 KRASi / DMSO (4 h)
B
Interferon Signaling
Cell Cycle & DNA Replication
DNA Damage Response
Extracellular Matrix
Integrins
Amino Acid Metabolism
Transcription
Fatty Acid and Lipid Metabolism
Smad Signaling
TCA Cycle
Lysosome
Translation
Glucose Metabolism
GSEA KRASi IC90 24 h vs. DMSO
C
mTOR signaling
Cholesterol synthesis
Cytoskeleton
Growth factor signaling
mRNA processing
Lipid metabolism and lysosome
Chondroitin sulfate/heparin
Cell cycle and translation
Protein folding
Immune response
Glutathione metabolism
Oxphos/ electron transport
Extracellular matrix
Nucleotide metabolism
Glucose metabolism
GSEA KRASi 8 w vs. DMSO
D
24 h top downregulated
Normalized enrichment score (NES)
Time
Class
Cell cycle
DNA repair
E
7 d top downregulated
Class
Actin cytoskeleton
Glucose metabolism and pentose phosphate
RNA metabolism
Translation and ribosome
F
24 h top upregulated
Class
ABC transporters
Aminoacid metabolism
Lipid metabolism
Lysosome
Perosixome
Propanoate and butanoate metabolism
TCA cycle and respiration
G
7 d top upregulated
Class
ECM and adhesion
Lysosome
Matrisome
Mitochondrial protein import
Sphingolipid metabolism
TCA cycle and respiration
H
Lysosome and mTOR
1h 4h 24h 72h 7d 8w

**Figure S2**

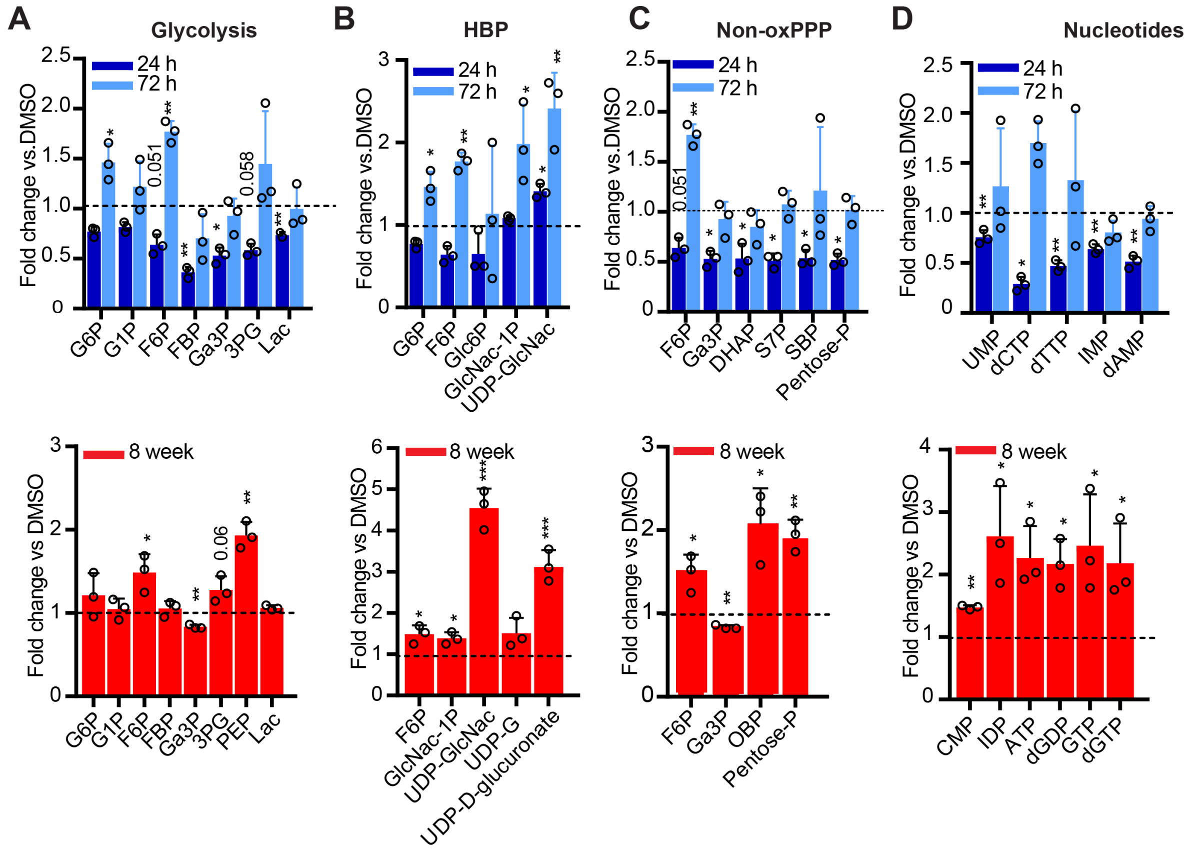

A
Glycolysis
B
HBP
C
Non-oxPPP
D
Nucleotides
24 h
72 h
8 week
Fold change vs.DMSO
Fold change vs DMSO
G6P
G1P
F6P
FBP
Ga3P
3PG
Lac
PEP
Glc6P
GlcNac-1P
UDP-GlcNac
UDP-G
UDP-D-glucuronate
DHAP
S7P
SBP
OBP
Pentose-P
UMP
dCTP
dTTP
IMP
dAMP
CMP
IDP
ATP
dGDP
GTP
dGTP
0.051
0.058
0.06

### Figure S3

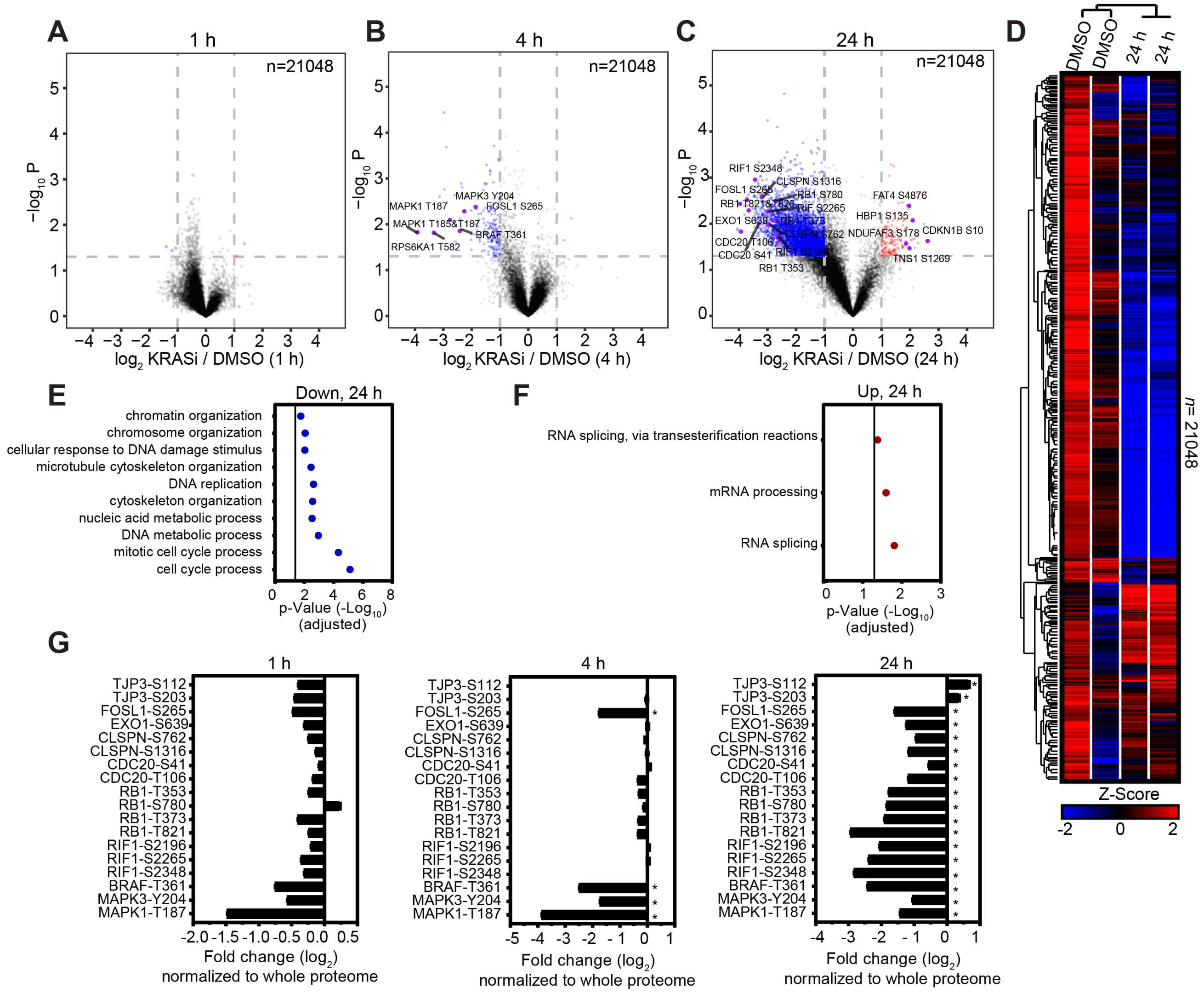

A
1 h
n=21048
−log10 P
log2 KRASi / DMSO (1 h)
B
4 h
n=21048
MAPK3 Y204
FOSL1 S265
MAPK1 T187
MAPK1 T185&T187
BRAF T361
RPS6KA1 T582
log2 KRASi / DMSO (4 h)
C
24 h
n=21048
RIF1 S2348
CLSPN S1316
FOSL1 S265
RB1 S780
RB1 T821&T826
RIF S2265
EXO1 S639
RB1 T373
CDC20 T106
CLSPN S762
CDC20 S41
RIF1 S2196
RB1 T353
FAT4 S4876
HBP1 S135
NDUFAF3 S178
CDKN1B S10
TNS1 S1269
log2 KRASi / DMSO (24 h)
D
DMSO
DMSO
24 h
24 h
n= 21048
Z-Score
-2
0
2
E
Down, 24 h
chromatin organization
chromosome organization
cellular response to DNA damage stimulus
microtubule cytoskeleton organization
DNA replication
cytoskeleton organization
nucleic acid metabolic process
DNA metabolic process
mitotic cell cycle process
cell cycle process
p-Value (-Log10)
(adjusted)
F
Up, 24 h
RNA splicing, via transesterification reactions
mRNA processing
RNA splicing
p-Value (-Log10)
(adjusted)
G
1 h
4 h
24 h
TJP3-S112
TJP3-S203
FOSL1-S265
EXO1-S639
CLSPN-S762
CLSPN-S1316
CDC20-S41
CDC20-T106
RB1-T353
RB1-S780
RB1-T373
RB1-T821
RIF1-S2196
RIF1-S2265
RIF1-S2348
BRAF-T361
MAPK3-Y204
MAPK1-T187
Fold change (log2)
normalized to whole proteome

#### Figure S4

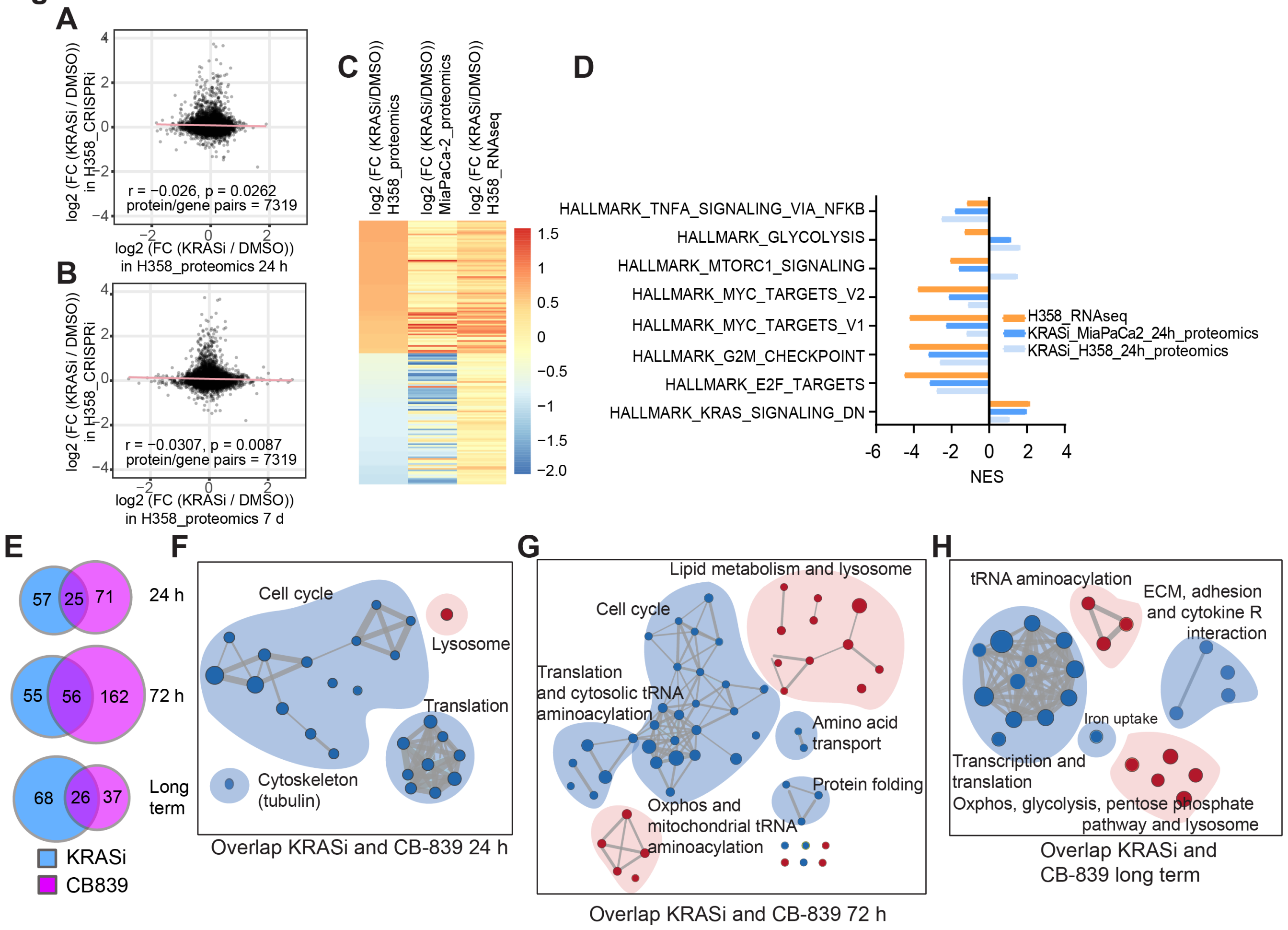

A
r = −0.026, p = 0.0262
protein/gene pairs = 7319
log2 (FC (KRASi / DMSO)) in H358_CRISPRi
log2 (FC (KRASi / DMSO)) in H358_proteomics 24 h
B
r = −0.0307, p = 0.0087
protein/gene pairs = 7319
log2 (FC (KRASi / DMSO)) in H358_CRISPRi
log2 (FC (KRASi / DMSO)) in H358_proteomics 7 d
C
log2 (FC (KRASi/DMSO)) H358_proteomics
log2 (FC (KRASi/DMSO)) MiaPaCa-2_proteomics
log2 (FC (KRASi/DMSO)) H358_RNAseq
D
HALLMARK_TNFA_SIGNALING_VIA_NFKB
HALLMARK_GLYCOLYSIS
HALLMARK_MTORC1_SIGNALING
HALLMARK_MYC_TARGETS_V2
HALLMARK_MYC_TARGETS_V1
HALLMARK_G2M_CHECKPOINT
HALLMARK_E2F_TARGETS
HALLMARK_KRAS_SIGNALING_DN
H358_RNAseq
KRASi_MiaPaCa2_24h_proteomics
KRASi_H358_24h_proteomics
NES
E
24 h
72 h
Long term
KRASi
CB839
F
Cell cycle
Lysosome
Translation
Cytoskeleton (tubulin)
Overlap KRASi and CB-839 24 h
G
Lipid metabolism and lysosome
Cell cycle
Translation and cytosolic tRNA aminoacylation
Amino acid transport
Protein folding
Oxphos and mitochondrial tRNA aminoacylation
Overlap KRASi and CB-839 72 h
H
tRNA aminoacylation
ECM, adhesion and cytokine R interaction
Iron uptake
Transcription and translation
Oxphos, glycolysis, pentose phosphate pathway and lysosome
Overlap KRASi and CB-839 long term

#### Figure S5

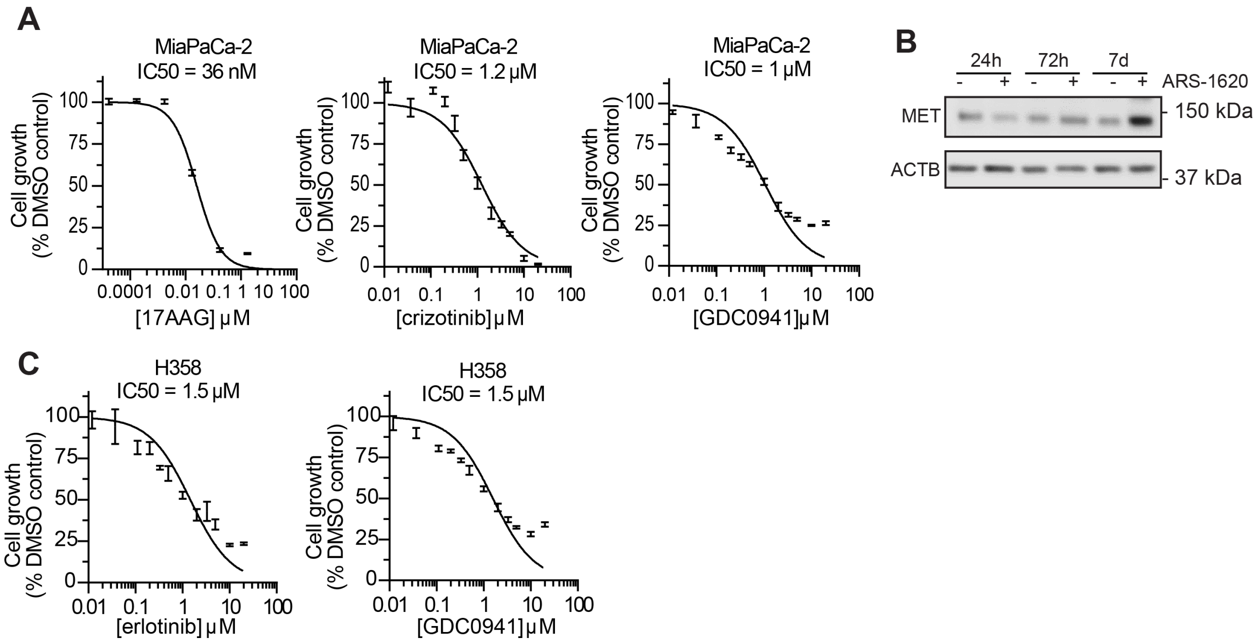
A
MiaPaCa-2
IC50 = 36 nM
Cell growth (% DMSO control)
[17AAG] μM
MiaPaCa-2
IC50 = 1.2 μM
[crizotinib] μM
MiaPaCa-2
IC50 = 1 μM
[GDC0941] μM
B
24h
72h
7d
ARS-1620
MET
ACTB
150 kDa
37 kDa
C
H358
IC50 = 1.5 μM
[erlotinib] μM
H358
IC50 = 1.5 μM
[GDC0941] μM

#### Figure S6

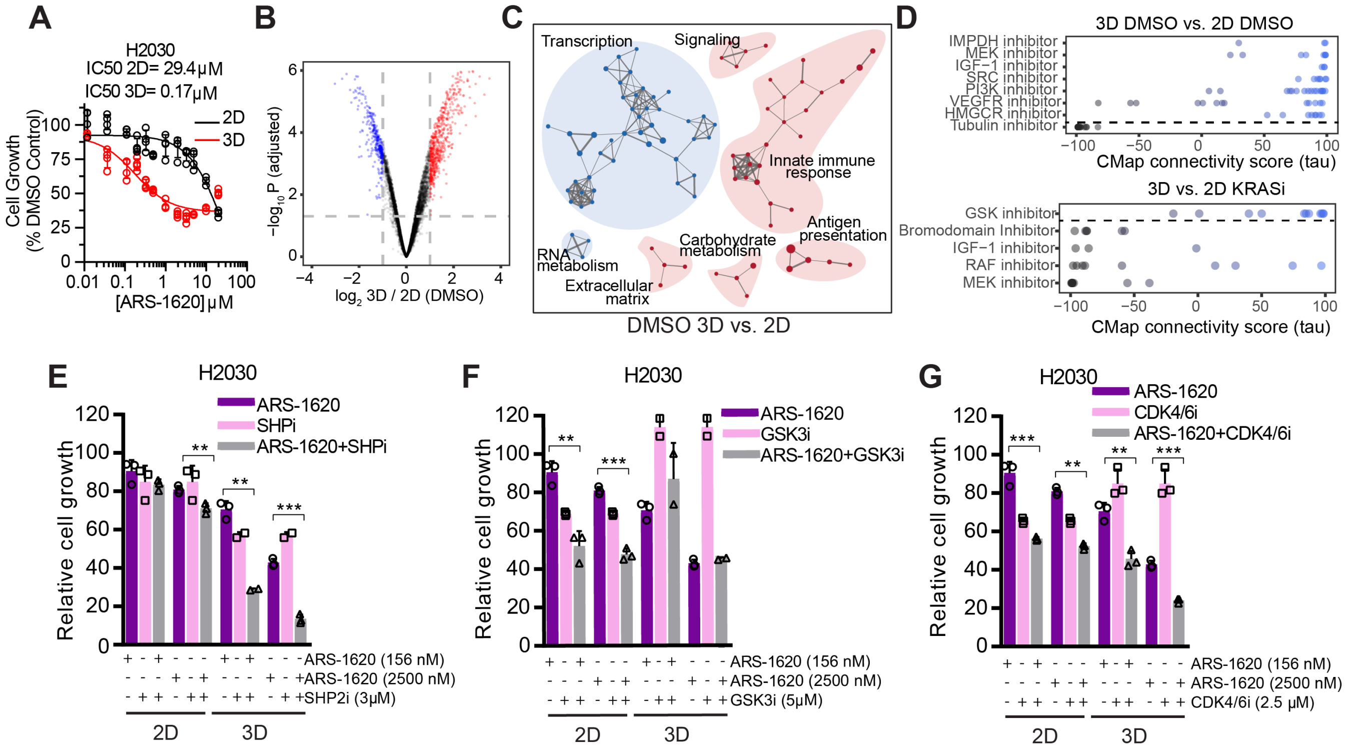

A
H2030
IC50 2D= 29.4μM
IC50 3D= 0.17μM
2D
3D
Cell Growth (% DMSO Control)
[ARS-1620] μM
B
−log10 P (adjusted)
log2 3D / 2D (DMSO)
C
Transcription
Signaling
Innate immune response
Antigen presentation
Carbohydrate metabolism
RNA metabolism
Extracellular matrix
DMSO 3D vs. 2D
D
3D DMSO vs. 2D DMSO
IMPDH inhibitor
MEK inhibitor
IGF−1 inhibitor
SRC inhibitor
PI3K inhibitor
VEGFR inhibitor
HMGCR inhibitor
Tubulin inhibitor
CMap connectivity score (tau)
3D vs. 2D KRASi
GSK inhibitor
Bromodomain Inhibitor
IGF−1 inhibitor
RAF inhibitor
MEK inhibitor
CMap connectivity score (tau)
E
H2030
ARS-1620
SHPi
ARS-1620+SHPi
Relative cell growth
ARS-1620 (156 nM)
ARS-1620 (2500 nM)
SHP2i (3μM)
2D
3D
F
H2030
ARS-1620
GSK3i
ARS-1620+GSK3i
Relative cell growth
ARS-1620 (156 nM)
ARS-1620 (2500 nM)
GSK3i (5μM)
2D
3D
G
H2030
ARS-1620
CDK4/6i
ARS-1620+CDK4/6i
Relative cell growth
ARS-1620 (156 nM)
ARS-1620 (2500 nM)
CDK4/6i (2.5 μM)
2D
3D

#### Figure S7

### A

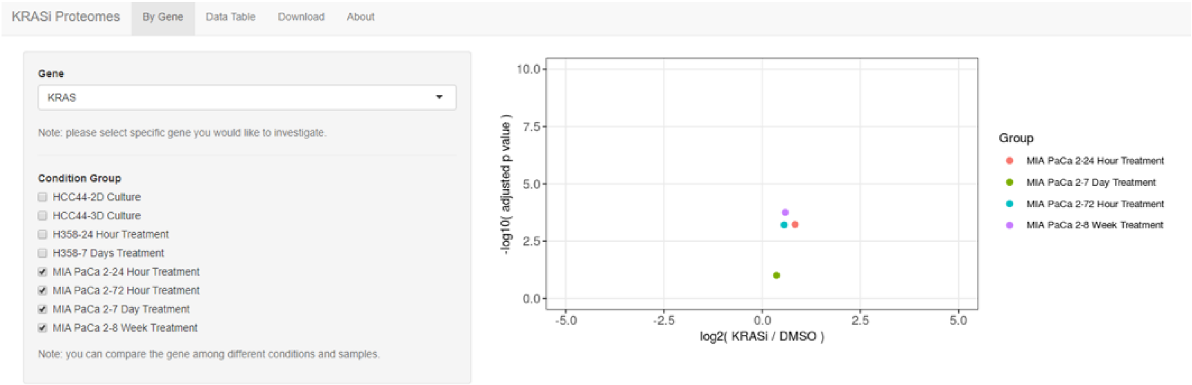

KRASi Proteomes
By Gene
Data Table
Download
About
Gene
KRAS
Note: please select specific gene you would like to investigate.
Condition Group
HCC44-2D Culture
HCC44-3D Culture
H358-24 Hour Treatment
H358-7 Days Treatment
MIA PaCa 2-24 Hour Treatment
MIA PaCa 2-72 Hour Treatment
MIA PaCa 2-7 Day Treatment
MIA PaCa 2-8 Week Treatment
Note: you can compare the gene among different conditions and samples.
-log10( adjusted p value )
log2( KRASi / DMSO )
Group
MIA PaCa 2-24 Hour Treatment
MIA PaCa 2-7 Day Treatment
MIA PaCa 2-72 Hour Treatment
MIA PaCa 2-8 Week Treatment

#### Figure S8

Full unedited immunoblot for Figure 1D: pAKT, AKT, pERK, ERK, KRAS, VINC

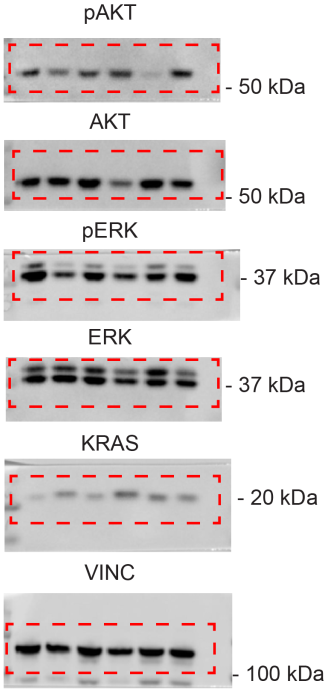


Full unedited immunoblot for Figure 2C: pAKT, AKT, pERK, ERK, KRAS, VINC

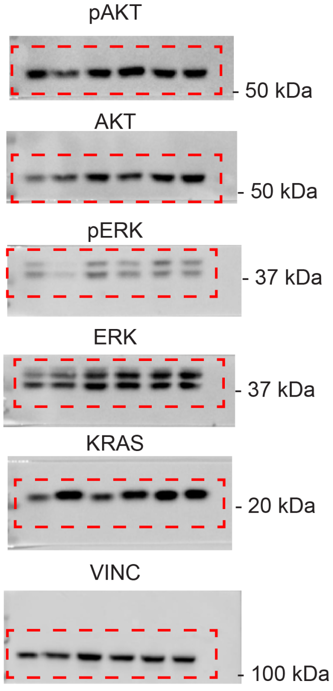


Full unedited immunoblot for Figure 6B: pAKT, AKT, pERK, ERK, KRAS, VINC

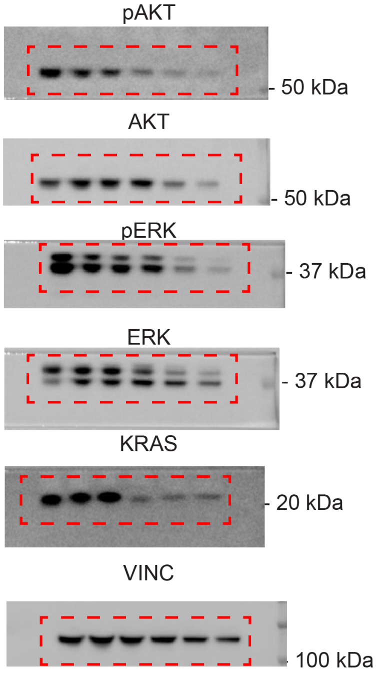


Full unedited immunoblot for Figure S5B: MET, ACTB

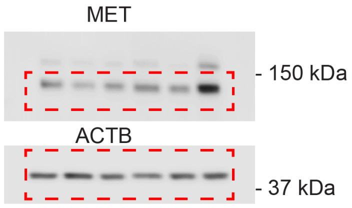
